# A rationally designed oral vaccine induces Immunoglobulin A in the murine gut that directs the evolution of attenuated *Salmonella* variants

**DOI:** 10.1101/824821

**Authors:** Médéric Diard, Erik Bakkeren, Verena Lentsch, Andrea Rocker, Nahimi Amare Bekele, Daniel Hoces, Selma Aslani, Markus Arnoldini, Flurina Böhi, Kathrin Schumann-Moor, Jozef Adamcik, Luca Piccoli, Antonio Lanzavecchia, Beth M. Stadtmueller, Nicholas Donohue, Marjan W. van der Woude, Alyson Hockenberry, Patrick H. Viollier, Laurent Falquet, Daniel Wüthrich, Ferdinando Bonfiglio, Claude Loverdo, Adrian Egli, Giorgia Zandomeneghi, Raffaele Mezzenga, Otto Holst, Beat H. Meier, Wolf-Dietrich Hardt, Emma Slack

**Affiliations:** Institute for Microbiology, Department of Biology, ETH Zürich, Zürich, Switzerland; Biozentrum, University of Basel, Basel, Switzerland; Institute for Food, Nutrition and Health, ETH Zürich, Zürich, Switzerland; Institute for Research in Biomedicine, Università della Svizzera italiana, Bellinzona, Switzerland; Department of Biochemistry, University of Illinois at Urbana-Champaign, Urbana, Illinois USA; York Biomedical Research Institute, Hull York Medical School, University of York, York, UK; Department of Environmental Microbiology, Eawag, Dubendorf, Switzerland; Department of Environmental Sciences, ETH Zürich, Switzerland; Microbiology and Molecular Medicine, University of Geneva, Geneva, Switzerland; Department of Biology, University of Fribourg, Fribourg, Switzerland; Swiss Institute of Bioinformatics, Fribourg, Switzerland; Infection Biology, Basel University Hospital, Basel, Switzerland; Department of Biomedicine, University of Basel, Basel, Switzerland; Institute for Physical Chemistry, ETH Zürich, Zürich, Switzerland; ETH Zürich, Department of Materials, Wolfgang-Pauli-Strasse 10, 8093 Zürich.; Forschungszentrum Borstel, Borstel, Germany

## Abstract

The ability of gut bacterial pathogens to escape immunity by antigenic variation, particularly via changes to surface-exposed antigens, is a major barrier to immune clearance^1^. However, not all variants are equally fit in all environments^2, 3^. It should therefore be possible to exploit such immune escape mechanisms to direct an evolutionary trade-off. Here we demonstrated this phenomenon using *Salmonella enterica* subspecies *enterica* serovar Typhimurium (*S.*Tm). A dominant surface antigen of *S.*Tm is its O-antigen: A long, repetitive glycan that can be rapidly varied by mutations in biosynthetic pathways or by phase-variation^4, 5^. We quantified the selective advantage of O-antigen variants in the presence and absence of O-antigen specific IgA and identified a set of evolutionary trajectories allowing immune escape without an associated fitness cost in naïve mice. Through the use of oral vaccines, we rationally induced IgA responses blocking all of these trajectories, which selected for *Salmonella* mutants carrying deletions of the O-antigen polymerase *wzyB.* Due to their short O-antigen, these evolved mutants were more susceptible to environmental stressors (detergents, complement), predation (bacteriophages), and were impaired in gut colonization and virulence in mice. Therefore, a rationally induced cocktail of intestinal antibodies can direct an evolutionary trade-off in *S.*Tm. This lays the foundations for the exploration of mucosal vaccines capable of setting evolutionary traps as a prophylactic strategy.

## Main text

The gut is a challenging environment for bacteria with high densities of phage, bile acids, antimicrobial peptides and secretory antibodies. These interact first with the outermost layer of the bacterial surface. Long, repetitive glycans, such as capsular polysaccharide, teichoic acids or O-antigens are ubiquitous as the outermost defense in bacteria. A particularly relevant feature of these glycan structures is that small changes in the structure of the repeating units, such as gain or loss of acetyl groups, when polymerized, result in major changes in conformation and charge-distribution of the glycans.

In the case of non-Typhoidal *Salmonella* this outermost glycan layer is predominantly made up of O-antigen: lipopolysaccharide core-linked, long, repetitive heteroglycans that hide most common outer-membrane proteins (^6, 7^, Fig.ED1). The *S.*Tm wild type (*S.*Tm^WT^) O:4[5], 12-0 O-antigen is a polymer of a triose repeating backbone (-mannose-α-(1→4)-rhamnose-α-(1→3)-galactose-α-(1→2), constituting the O:12-0 epitope) with an α-(1→3)-abequose side-branch at the mannose (constituting the O:4 epitope, or when *O*-acetylated the O:5 epitope) (Fig. 1A). The *S.*Tm^WT^ reacts to O:5-typing antisera and O:12-0-typing antibodies (Fig. 1B, **and** C, S1-3). In the SL1344 strain of *S.*Tm, two major shifts in O-antigen composition have been reported. Firstly, complete loss of abequose acetylation, generating an O:4-only phenotype, occurs via loss of function mutations in the abequose acetyl transferase gene *oafA*^8^, (Fig. 1A and B). Secondly, the O:12-0 epitope can be converted to an O:12-2 epitope by (α-(1→4) glucosylation of the backbone galactose (Fig. 1A and C). This occurs via expression of a glucosyl transferase *gtrABC* operon (STM0557-0559), controlled by DAM-dependent methylation i.e. by phase variation^4, 9^. Note that *S*.Tm strain SL1344 lacks a second common operon required for linking glucose via an α-(1→6) linkage to the backbone galactose, generating the O:1 serotype. All of these structural O-antigen variants exert only a mild fitness defect in the naïve gut (^5, 9, 10^, Fig 1D and E). However, there is also evidence for selection of mutants at loci coding for the O-antigen polymerases and so-called “non-typable” *Salmonella* strains with a single-repeat O-antigen are occasionally observed amongst isolates from infected humans or animals^11^. Such strains lose outer membrane robustness, due to loss of the rigid hydrophilic glycan layer^12^,. and therefore have decreased fitness both in the gut and in the environment^2, 3, 13^.

**Figure 1:**
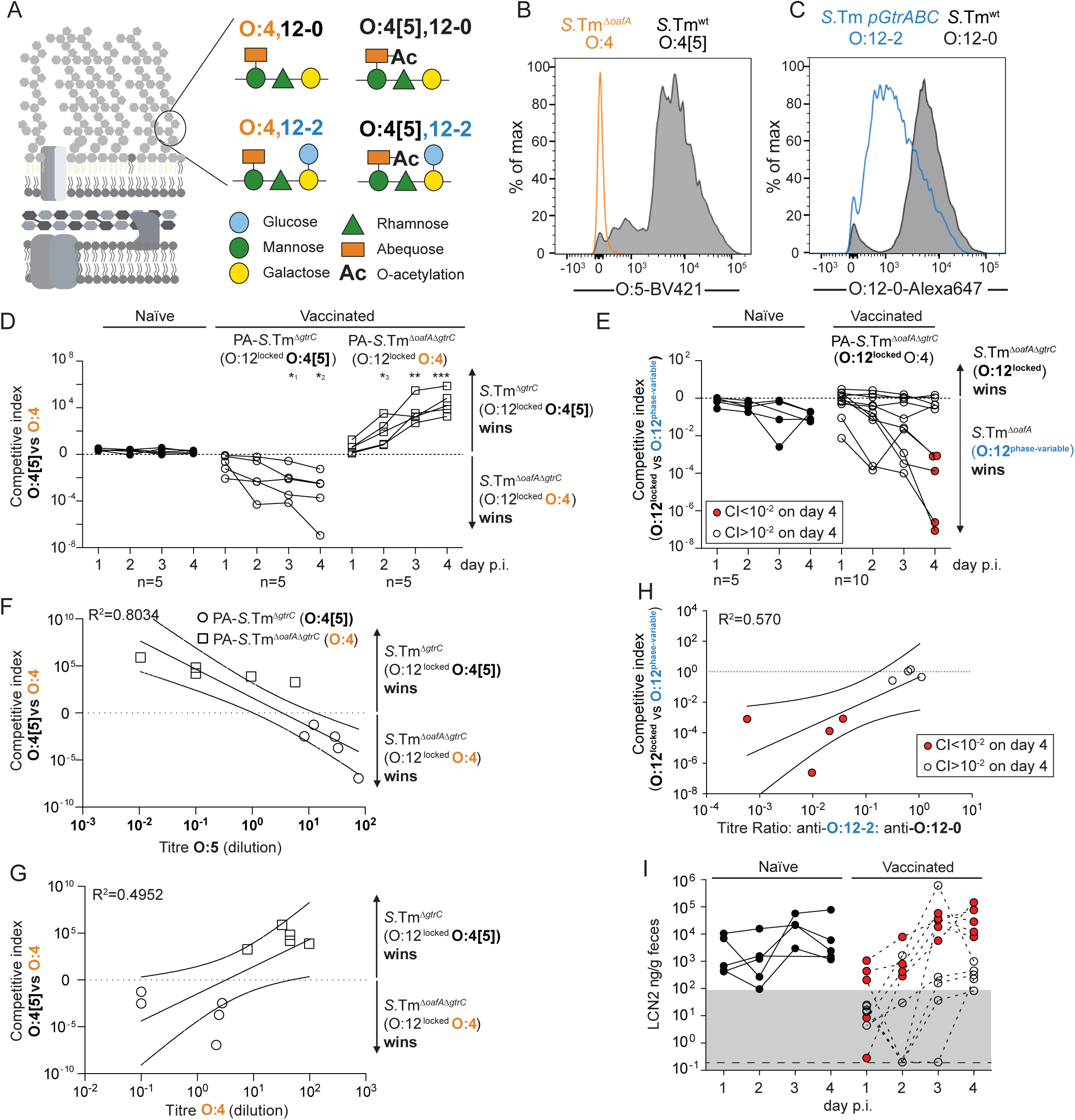
Vaccine-induced IgA exerts a strong selective pressure on O-antigen variants during murine non-Typhoidal *Salmonellosis*: **A**. Schematic of the O-antigen of *S*.Tm (O:4[5],12), and its common variants depicted using the “Symbol Nomenclature for Glycans”. **B and C**. Overnight cultures of the indicated *S.*Tm strains were stained for presence of O:5 **(B)** or O:12-0 **(C)** epitopes. **(D-I)** Naïve and vaccinated C57BL/6 mice were streptomycin-pretreated and infected with the indicated combination of *S.*Tm strains. **(D,F,G)** Naïve (closed circles, n=5), PA-S.Tm^Δ*gtrC*^-vaccinated (O:4[5]-vaccinated, open circles, n=5) and PA-S.Tm^Δ*gtrC*Δ*oafA*^-vaccinated (O:4-vaccinated, open squares, n=5) SPF mice were streptomycin-pretreated, infected (10^5^ CFU, 1:1 ratio of *S.*Tm^Δ*gtrC*^ and *S.*Tm^Δ*gtrC* Δ*oafA*^ per os). **D**. Competitive index (CFU *S.*Tm ^Δ*gtrC*^*/*CFU *S.*Tm^Δ *gtrC* Δ*oafA*^) in feces at the indicated time-points. Two-way ANOVA with Bonferroni post-tests on log-normalized values, compared to naive mice. *^1^p=0.0443, *^2^p=0.0257, *^1^p=0.0477, **p=0.0021,***p=0.0009 **F** and **G**. Correlation of the competitive index with the O:4[5]-binding (**F**) and O:4-binding (**G**) intestinal IgA titre, r^2^ values of the linear regression of log-normalized values. Open circles: Intestinal IgA from O:4[5]-vaccinated mice, Open squares: Intestinal IgA from O:4-vaccinated mice. Lines indicate the best fit with 95% confidence interval. **E,H, I.** Naive (closed circles, n=5) or PA-S.Tm ^Δ*oafA* Δ*gtrC*^ -vaccinated (O:4/O:12-0-vaccinated, open circles and red circles, n=10) C57BL/6 mice were streptomycin-pretreated and infected (10^5^ CFU, 1:1 ratio of *S.*Tm^Δ*oafA*^ (O:12-2 switching) and *S.*Tm ^Δ*oafA* Δ*gtrC*^ (O:12-locked) *per os*). **E**. Competitive index (CFU *S.*Tm^Δ*oafA* Δ*gtrC*^ */*CFU *S.*Tm^Δ*oafA*^) in feces at the indicated time-points. Red circles indicate vaccinated mice with a competitive index below 10^-2^ on d4 and are used to identify these animals in panels **H and I**. **E** Effect of vaccination is not significant by 2-way ANOVA considering vaccination over time. **H**. Correlation of the competitive index on day 4 with the ratio of intestinal IgA titre against an O:12-2-locked *S.*Tm p*gtrABC* variant to the titre again an O:12-0-locked *S.*Tm*^GtrC^* variant (linear regression of log-normalized values, lines indicate the best fit with 95% confidence interval). **I**. Intestinal inflammation, corresponding to mice in panel E, quantified by measuring Fecal Lipocalin 2 (LCN2).

We hypothesized that the host’s immune response could generate conditions in which the fitness of O-antigen polymerase mutants is promoted, driving the emergence of an evolutionary trade-off. Intestinal antibodies (predominantly secretory IgA) are known to exert specific selective pressures on targeted species^14–16^. In order to investigate the evolutionary consequences of vaccine-induced secretory antibody responses in the gut, without the major ecological shifts associated with live-attenuated vaccine infection^17–19^, we made use of an established high dose, inactivated oral vaccination technique^15, 20, 21^ that induces intestinal IgA responses without detectable intestinal damage, inflammation or colonization by the vaccine strains^21^. Our standard vaccine (“PA-S.Tm”) consists of concentrated peracetic acid killed bacteria^21^. Conventional mice harboring a complex microbiota (16S amplicon analysis available^22^) received 10^10^ particles of PA-S.Tm orally once per week for 4 weeks. Subsequently, these mice were antibiotic-treated to open a niche for the pathogen in the large intestine, and were infected with *S.*Tm SL1344, which rapidly colonizes the cecum, generating typhlocolitis, and invasive disease in the mesenteric lymph nodes, spleen and liver^23, 24^.

We first quantified the competitive fitness of *S.*Tm mutants genetically “locked” into individual structural O-antigen compositions in vaccinated and naïve mice. Competition between *S.*Tm^Δ*oafA* Δ*gtrC*^ (**O:4**, O:12-0-locked) and *S.*Tm^Δ*gtrC*^ (**O:4[5]**, O:12-0-locked) demonstrated no difference in fitness in naïve mice over 4 days of infection. However, in mice vaccinated either against the O:4 or the O:4[5] variant (Fig. S4), we observed up to a 10^7^-fold outcompetition of the IgA-targeted O-antigen variant within 4 days (Fig. 1D). The magnitude of the selective advantage correlated with the magnitude of the intestinal IgA response to each O-antigen variant (Fig. 1F and G). Therefore, IgA can exert a strong selective pressure on the O:4/O:4[5] O-antigen variants. Competing *S.*Tm^Δ*oafA*^ (**O:12-phase-variable,** O:4) against *S.*Tm^Δ*oafA* Δ*gtrC*^ (**O:12-locked,** O:4) revealed a mild benefit of O:12 phase variation in naïve mice up to day 4 post-infection, in line with published data (Fig. 1E) ^4, 5^. However, we observe a major fitness benefit of phase variation in vaccinated mice in which the IgA response is highly biased to recognition of O:12-0 O-antigens (Fig. 1E, H. Red symbols, Fig. S5). Correspondingly, vaccinated mice with an outgrowth of phase-variable *S.*Tm also displayed initiation of intestinal inflammation, as quantified by fecal Lipocalin 2 (LCN2, Fig. 1I). The mechanistic basis of this selective advantage could be confirmed by complementation of the *gtrC* gene in trans (Fig. S6). Therefore O:12-0-targeting IgA can exert a strong selective pressure against *S.*Tm unable to phase-vary the O:12-0 part of the O-antigen. As neither of these variants (O:4[5] to O:4 and O:12-0 to O:12-2) are associated with a major loss-of-fitness in naïve mice (Fig. 1D and E), this implied that such variants should be selected for during infections of vaccinated mice with wild type *Salmonella*.

We therefore established whether natural emergence of these “IgA-escape”-*S.*Tm variants occurred sufficiently fast to be observed during wild type *S.*Tm infections. For this purpose, we treated mice with a wild type PA-S.Tm oral vaccine as above, or with a vehicle-only control, and then challenged these animals with wild type *S.*Tm. Around 30% of vaccinated mice showed intestinal inflammation at 18 h post infection (Fig. 2A), despite the presence of robust anti-*S.*Tm^wt^ intestinal IgA in all vaccinated animals (Fig. 2B). When *S.*Tm clones were recovered from the cecal content of vaccinated mice with intestinal inflammation, these were typically recognized less well by vaccine-induced IgA than *S.*Tm clones from the cecum of vaccinated and protected mice (Fig. 2C). In 11 of 34 mice analysed, we observed clones with complete loss-of-binding to an O:5-specific polyclonal antisera within 4 days (Table S3, Fig 2D). Resequencing of O:5-negative clones confirmed a 7 bp contraction of a tandem repeat in the open reading frame of *oafA,* coding for the abequose acetylase (Fig. 2E, 10 different clones from three independent experiments), that is also found in multiple NCBI deposited genomes^25^ (Fig. ED2A). A second site of microsatellite instability is present in the promoter of *oafA* suggesting a further possibility for rapid inactivation (Fig. ED2B), and this gene was found to be under negative selection in a recent screen of published *Salmonella* genomes^26^.

**Figure 2:**
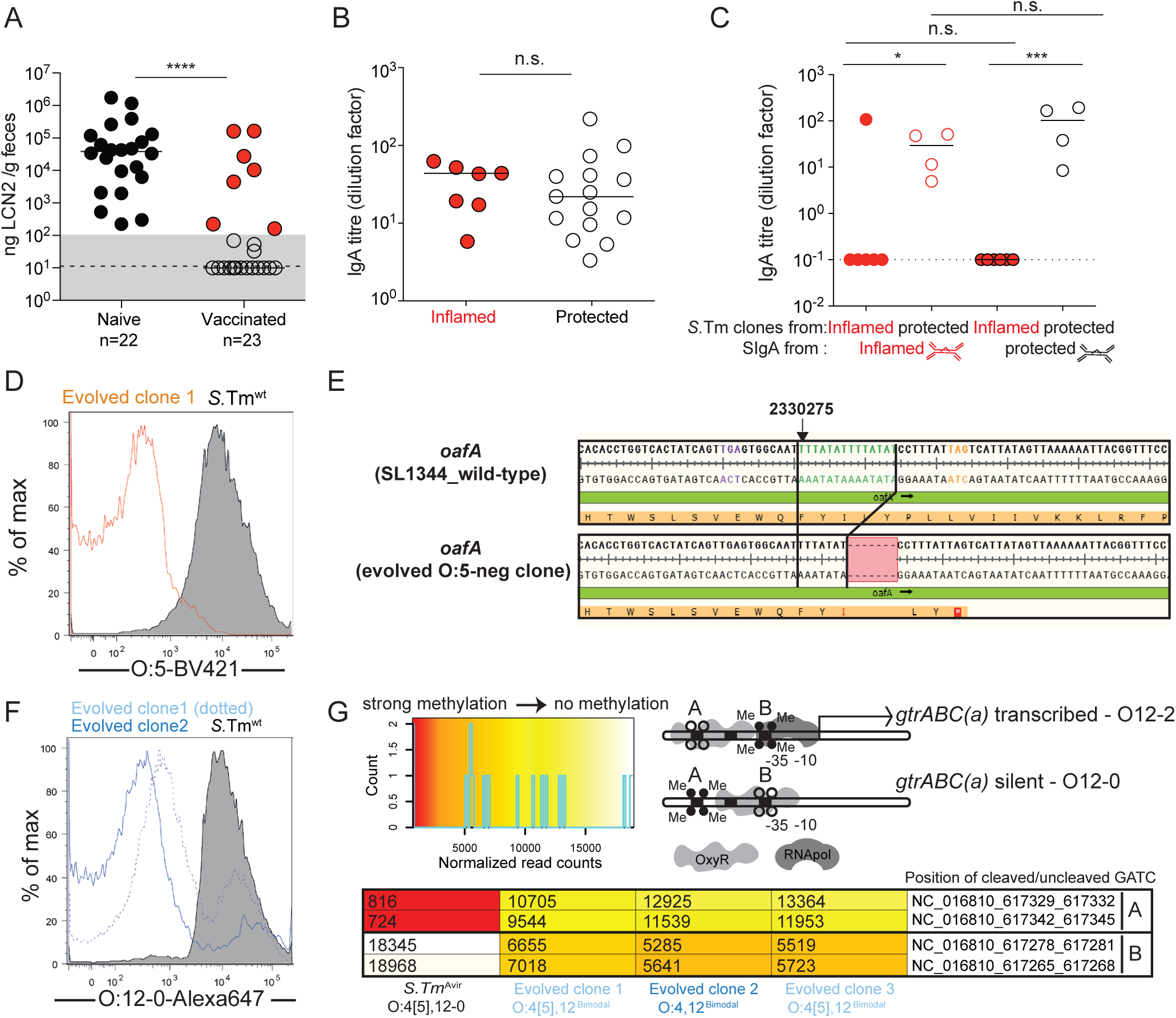
O-Antigen variants rapidly emerge during wild type *S.*Tm infection of vaccinated mice: **A-C** : Naïve (n=22) or PA-S.Tm-vaccinated (Vaccinated, n=23) SPF C57BL/6 mice were streptomycin-pretreated, infected (10^5^ *S.*Tm^wt^ Colony forming units (CFU) per os) and analyzed 18 h later. **A**. Fecal Lipocalin 2 (LCN2) to quantify intestinal inflammation, 2-tailed Mann Whitney U test p<0.0001 **B**. Intestinal IgA titres against *S*.Tm^wt^ determined by flow cytometry, for vaccinated mice with LCN2 values below (open symbols, protected) and above (filled symbols, inflamed) 100ng/g. p=0.61 by 2-tailed Mann Whitney U test. **C.** Titres of intestinal lavage IgA from an “inflamed vaccinated” mouse (red borders) or a “protected vaccinated” mouse (black borders) against *S.*Tm clones re-isolated from the feces of the “inflamed vaccinated” mouse (red filled circles) or “protected vaccinated” mouse (open circles) at day 3 post-infection. Two-way ANOVA with Bonferroni post-tests on log-normalized data. Clones and lavages from n=1 mouse, representative of 9 “vaccinated but inflamed” and 13 “vaccinated protected” mice, summarized in Table S4. *p=0.0156, ***p=0.0003. **D.** Flow cytometry staining of *S.*Tm^wt^ and an evolved with anti-O:5 typing sera (gating as in Fig. S1). **E.** Alignment of the *oafA* sequence from wild type (SL1344_RS11465) and an example O:5-negative evolved clone showing the 7bp contraction leading to premature stop codon (all four re-sequenced O:5-negative strains showed the same deletion). **F.** Binding of an O:12-0-specific monoclonal antibody to *S.*Tm^wt^ and O:12^Bimodal^ evolved clones, determined by bacterial flow cytometry. (gating as in Fig. S1). **G.** Methylation status of the g*trABC* promoter region in *S.*Tm, and three O:12^Bimodal^ evolved clones determined by REC-seq. Heat-scale for normalized read-counts, schematic diagram of promoter methylation associated with ON and OFF phenotypes, and normalized methylation read counts for the indicated strains.

In contrast, loss of O:12-0 staining was bimodal within individual clones (Fig. 2F), consistent with phase-variation^4^ and no reproducible mutations were identified in these clones on genome resequencing (**Table S3**). Instead, methylation analysis revealed a methylation pattern indicative of the *gtrABC* promoter being in an “ON” conformation (Fig. 2G). Serial passage of these clones (Fig. ED3A), as well as cultivation in microfluidic devices (**Supplementary videos 1 and 2**) confirmed the ability of clones to switch between O:12-0-positive and negative states. The STM0557-0559 *gtrABC* locus was confirmed to be essential for this observed loss of O:12-0 epitope as strains lacking *gtrC* remained 100% O:12-0-positive even under strong *in vivo* selection (Fig. ED3B and C). This phenotype could be replicated by adoptive transfer of a recombinant monoclonal IgA specific for the O:12-0 epitope (mSTA121, Fig. ED4), confirming that O:12-0-binding IgA is sufficient to drive outgrowth of O:12-2-producing variants. Computational modeling of phase-variation and growth, as well as comparison of O:12-0/O12:2 switching rates of *lacZ* reporter strains suggested that selection for clones expressing *gtrABC* is sufficient to explain the recovery rate, without any intrinsic shift in phase variation switching rates (Fig. ED5). The chemical structure of O-antigen of the recovered clones was further confirmed by ^1^H-NMR of purified O-antigen and by high resolution magic-angle spinning NMR of O-antigen on the surface of intact cells (Fig. ED6). Therefore, vaccine-induced IgA can select for the natural emergence of O-antigen variants within a few days of infection with *S.*Tm wild type, resulting in disease in vaccinated mice. This phenomenon can also be observed at later time-points in IgA-competent but not IgA-deficient mice during chronic infection with live-attenuated *S.*Tm strains (Fig. ED7A and B) i.e. IgA is necessary for selection of O-antigen variants during chronic infection. Correspondingly, although the inactivated oral vaccines induce a higher titre of *Salmonella-*binding IgA than the live vaccines (Fig. ED7C), the response to chronic infection binds to O:4 and O4[5]-producing *S.*Tm with similar titres, while the response to inactivated vaccine is highly biased for the O-antigen variant of the vaccine (Fig. ED7C). This indicates that within-host O-antigen variation also occurs under the selective pressure of intestinal antibodies during chronic infections, and sequential priming will include a broad IgA response capable of recognizing multiple O-antigen variants.

We next investigated whether the relative fitness defect of a short O-antigen mutant can be compensated for by the selective advantage from lower IgA-binding in the gut lumen, i.e. whether IgA could drive an evolutionary trade-off. One-on-one competitions were carried out between *S.*Tm^Δ*oafA* Δ*gtrC*^ (O:4,12-0-locked**, long O-antigen**) and *S.*Tm^Δ*oafA* Δ*gtrC* Δ*wzyB*^ (O:4,12-0-locked**, short O-antigen,** retains just a single O-antigen repeat) in the intestine of mice with and without IgA raised against *S.*Tm^Δ*oafA* Δ*gtrC*^ (Fig. 3A). The single repeat O-antigen strain was rapidly outcompeted in naive animals, in line with earlier studies^11, 27^ (Fig. 3A) indicating a major loss-of-fitness. However, in the gut of vaccinated mice, strains with short O-antigen were dominant by day 4 (Fig. 3A). Vaccinated antibody-deficient mice were indistinguishable from naive mice in these experiments, verifying that IgA is necessary for the selection of short O-antigen strains in the gut of vaccinated mice (Fig. 3A). Introduction of day 4 fecal bacteria from vaccinated mice into naïve mice resulted in re-outgrowth of the strain with a long O-antigen, indicating that vaccine-induced IgA, and not secondary mutations in *S*.Tm^Δ*oafA* Δ*gtrC* Δ*wzyB*^, was responsible for competition outcome (Fig. 3B). The IgA titre recognizing short O-antigen-producing strains was lower than that against full-length O-antigen strains, consistent with the selective advantage in vaccinated mice (Fig. 3C). As the long O-antigen can have several hundred repeats of the glycan, decreased antibody binding could be driven by lower O-antigen abundance or by loss of avidity-driven interactions. Loss of long O-antigen can therefore be an advantage to *Salmonella* in the gut lumen of vaccinated mice.

**Figure 3:**
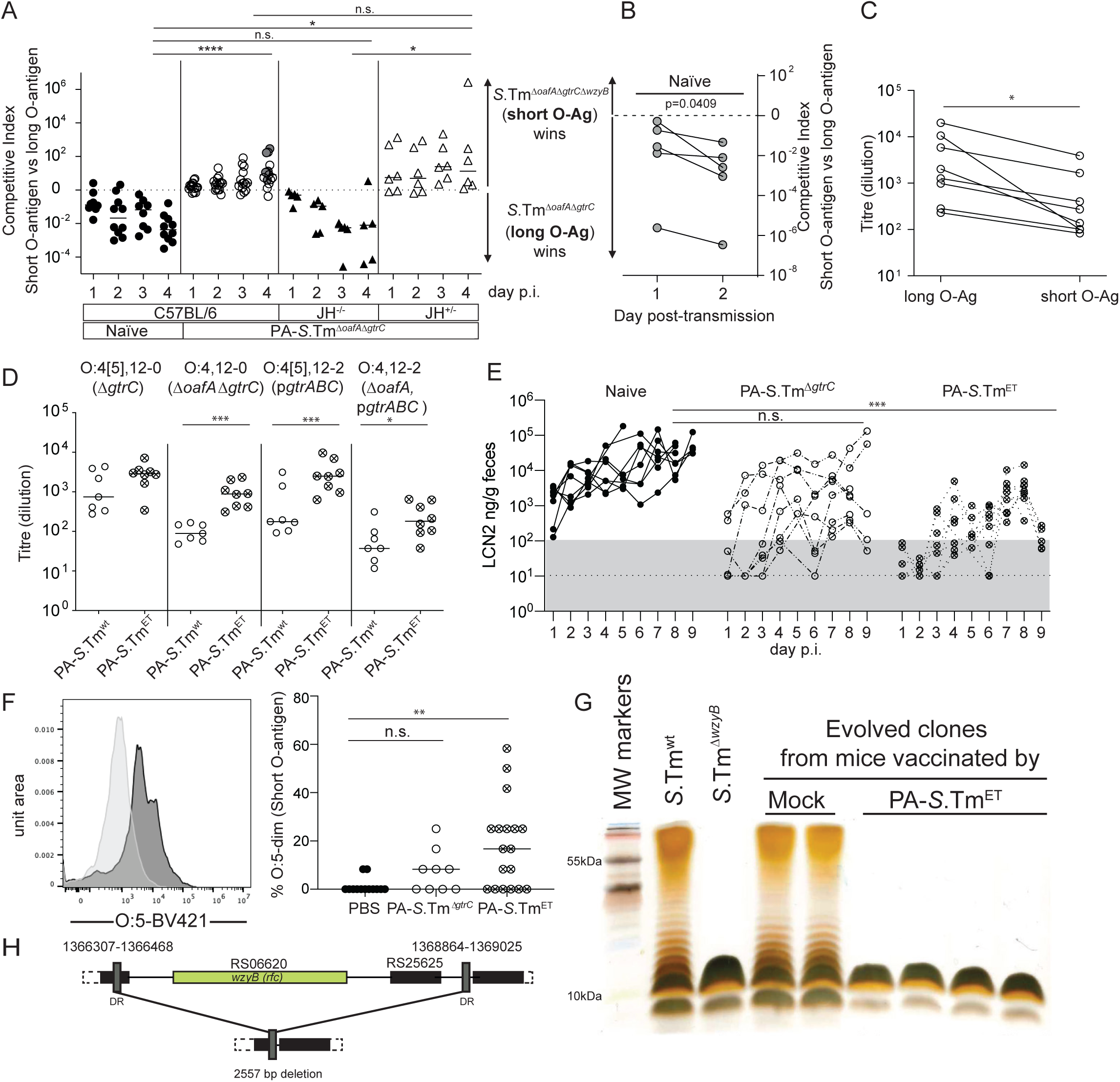
Single-repeat O-antigen confers a selective advantage in the presence of broad-specificity vaccine-induced IgA: **A-C**. Mock-vaccinated wild type (C57BL/6, n=10), PA-S.Tm^Δ*oafA* Δ*gtrC*^ *-*vaccinated JH^-/-^ mice (JH^-/-^n=6), PA-S.Tm^Δ*oafA* Δ*gtrC*^ *-*vaccinated wild type (C57BL/6, n=16) and PA-S.Tm^Δ*oafA* Δ*gtrC*^*-*vaccinated JH^+/-^ littermate controls (JH^+/-^, n=5 mice) were streptomycin pre-treated and infected with 10^5^ CFU of a 1:1 ratio *S.*Tm ^Δ*oafA* Δ*gtrC* Δ*wzyB*^ and *S.*Tm^Δ*oafA* Δ*gtrC*^ i.e. serotype-locked, short and long O-antigen-producing strains. **A.** Competitive index of *S.*Tm in feces on the indicated days. 2-way ANOVA with Tukey’s multiple comparisons tests. *p=0.0392, ****p<0.0001. **B**. Feces from the indicated mice (grey-filled circles panel **A)** were transferred into streptomycin-pretreated C57BL/6 naive mice (one fecal pellet per mouse, n=5). Competitive index in feces over 2 days of infection. **C.** Intestinal IgA titre from PA-S.Tm^Δ*oafA* Δ*gtrC*^ -vaccinated mice binding to *S.*Tm ^Δ*oafA* Δ*gtrC*^ (long O-antigen) and *S.*Tm^Δ*oafA* Δ*gtrC* Δ*wzyB*^ (short O-antigen). *p=0.0078 by 2-tailed Wilcoxon matched-pairs signed rank test. **D.** Intestinal IgA titre induced by PA-S.Tm^wt^ or PA-S.Tm^ET^ (4-strains) in 129S1/SvImJ mice determined by bacterial flow cytometry. Two-way ANOVA with Bonferroni multiple comparisons tests. Adjusted p values *p=0.0332, ***p=0007. (Gating Fig.S5, further data Fig. S7 and S8) **E.** 129S1/SvImJ Mice were vaccinated with vehicle only (Naïve, n=8), PA-S.Tm^wt^ (n=8), PA-S.Tm^ET^ (n=8). On day 28 after the first vaccination, mice were streptomycin pre-treated and challenged with 10^5^ *S.*Tm^wt^ orally. Intestinal inflammation as scored by fecal Lipocalin-2 (LCN2) days 1-9 post-infection. Dotted line = detection limit. Grey box = normal range in healthy mice. 2-way repeat-measures ANOVA with Tukey’s multiple comparison test. *** adjusted p value=0.0002 **F.** Representative plot of O:5 staining in an evolved clone with short O-antigen and quantification of the percentage of O:5-dim *S.*Tm clones re-isolated from the feces of infected SPF mice vaccinated with PBS only (n=13), PA-S.Tm^Δ*gtrC*^ (n=9) or PA-S.Tm^ET^ (n=18). Kruskal-Wallis test with Dunn’s multiple comparison tests shown. **p=0.0016. (gating as Fig. S1) **G.** Silver-stained gel of LPS from representative control and evolved *S.*Tm strains from 2 different control and vaccinated PA-S.Tm^ET^ mice. **H.** Resequencing of short O-antigen strains revealed a deletion between inverted repeats (n=5 clones, isolated from 2 different mice).

Based on these above observations, we hypothesized that emergence of mutants with a short O-antigen could be achieved for a wild type *S.*Tm infection if we could block all other IgA escape routes, effectively generating an evolutionary trap. To this end, mice received an oligovalent vaccine containing the **O:4[5],12** *S.*Tm^Δ*gtrC,*^, **O:4,12** *S.*Tm^Δ*oafA* Δ*gtrC*^, **O:4,12-2** *S.*Tm^Δ*oafA*^ p*gtrABC*, and **O:4[5],12-2** *S.*Tm p*gtrABC* strains (referred to as PA-*S*.Tm^ET^). This induced a broad antibody response with high avidity for all four of the known long O-antigen variants present in our *S.*Tm SL1344 strain (Fig. 3D, Fig.S7-8). PA-*S*.Tm^ET^ provided subtly better protection from intestinal inflammation in long-term infection of 129S1/SvImJ mice than the monovalent **O:5,12-0** vaccine (Fig. 3E, significant protection from intestinal inflammation at d9 with PA-*S*.Tm^ET^ but not PA-*S*.Tm^Δ*gtrC*^), as well as on mixed challenge of Balb/c mice (Fig. S9, significant protection from intestinal inflammation at d4 with PA-*S*.Tm^ET^ but not PA-*S*.Tm^Δ*gtrC*^). Moreover, our hypothesis that this vaccine can set an evolutionary trap was supported: short O-antigen-producing clones were detected in 12 of 18 PA-S.Tm^ET^ vaccinated mice analysed across multiple experiments by phenotypic characterization (anti-O5^dim^ flow cytometry staining, Fig. 3F). The O-antigen phenotype was confirmed by gel electrophoresis of purified LPS (Fig. 3G). Sequencing of evolved short-O-antigen clones (**Table S4**, n=5) revealed a common large deletion encompassing the *wzyB* gene (also termed *rfc*), encoding the O-antigen polymerase^11^ (Fig. 3H, Fig. ED8 also reported in some “non-typable” *S.*Tm isolates from broilers^11^). This deletion is mediated by site-specific recombination between flanking direct repeats, which renders the *wzyB* locus unstable^11^.

We have previously published that IgA responses against the surface of rough *Salmonella* are identically induced by vaccination with either rough or wild type *Salmonella* oral vaccines^28^. Correspondingly, including a short-O-antigen mutant into our PA-S.Tm^ET^ mix does not further improve IgA titres (Fig. S10). Note that in these experiments, we also do not observe a significant improvement of protection with PA-S.Tm^ET^, as PA-STm^WT^ protected well out to day 3 in n=6 of 8 mice, when the experiment was terminated for ethical reasons relating to the control group. As the generation of *Salmonella* O-antigen variants is inherently stochastic, but a prerequisite for selection by IgA and therefore within-host evolution, perfect protection can be observed in a variable fraction of animals that had received the monovalent vaccine up to this time-point. However, no intestinal inflammation, as quantified by fecal Lipocalin-2, was observed in any of the mice receiving PA-S.Tm^ET^ (n=9) or PA-S.Tm^ET+*wzyB*^ (n=4).

We finally confirmed that re-isolated *wzyB-*deletion mutants phenocopied the fitness defects of targeted *wzyB* mutations in harsh environments. Single infections with *S*.Tm^Δ*oafA* Δ*gtrC* Δ*wzyB*^ revealed that, in comparison to isogenic wild type counterparts, *wzyB*-deficient mutants (synthetic or evolved) are significantly less efficient at colonizing the gut of streptomycin pretreated naïve mice (Fig. 4A), disseminating systemically (Fig. 4B) and triggering inflammation (Fig. 4C), i.e. they have an intrinsic defect in colonization and virulence. This attenuation can be attributed to compromised outer membrane integrity^12^ and also manifests as an increased sensitivity to membrane destabilization by EDTA, bile acids and weak detergents (Fig ED9A-E) and increased sensitivity to complement-mediated lysis^11, 27^ (Fig. ED9F). It is also well-documented that specific interactions between the tail spike fiber and O-antigen reduce the host-range of ubiquitous lytic phages^29, 30^. Correspondingly, infection of the short-O-antigen strains with filtered wastewater generated visible lysis plaques of various sizes (Fig. 4D and E, Fig. ED10A). About 10-fold less lysis plaques were visible in the same conditions with long O-antigen strains (Fig. 4D and E, Fig. ED10A). Sequencing of phages isolated from four plaques revealed four different T5-like phages (Fig. 4F). Infections with the purified phage <u>φ12</u> yielded more phages after infection of a short O-antigen evolved clone compared to the ancestor strains (Fig. 4G). We could confirm that infection was dependent on *btuB*, the vitamin B12 outer-membrane transporter that is normally shielded by a long O-antigen (Fig, ED10B **and** C).These results confirmed that the recovered *wzyB* mutants were indeed sensitive to diverse membrane stresses, innate immune defenses and common environmental phages that would be encountered during transmission or on infection of a new host. Therefore, vaccination can successfully drive evolution toward fitness trade-off *in vivo*.

**Figure 4:**
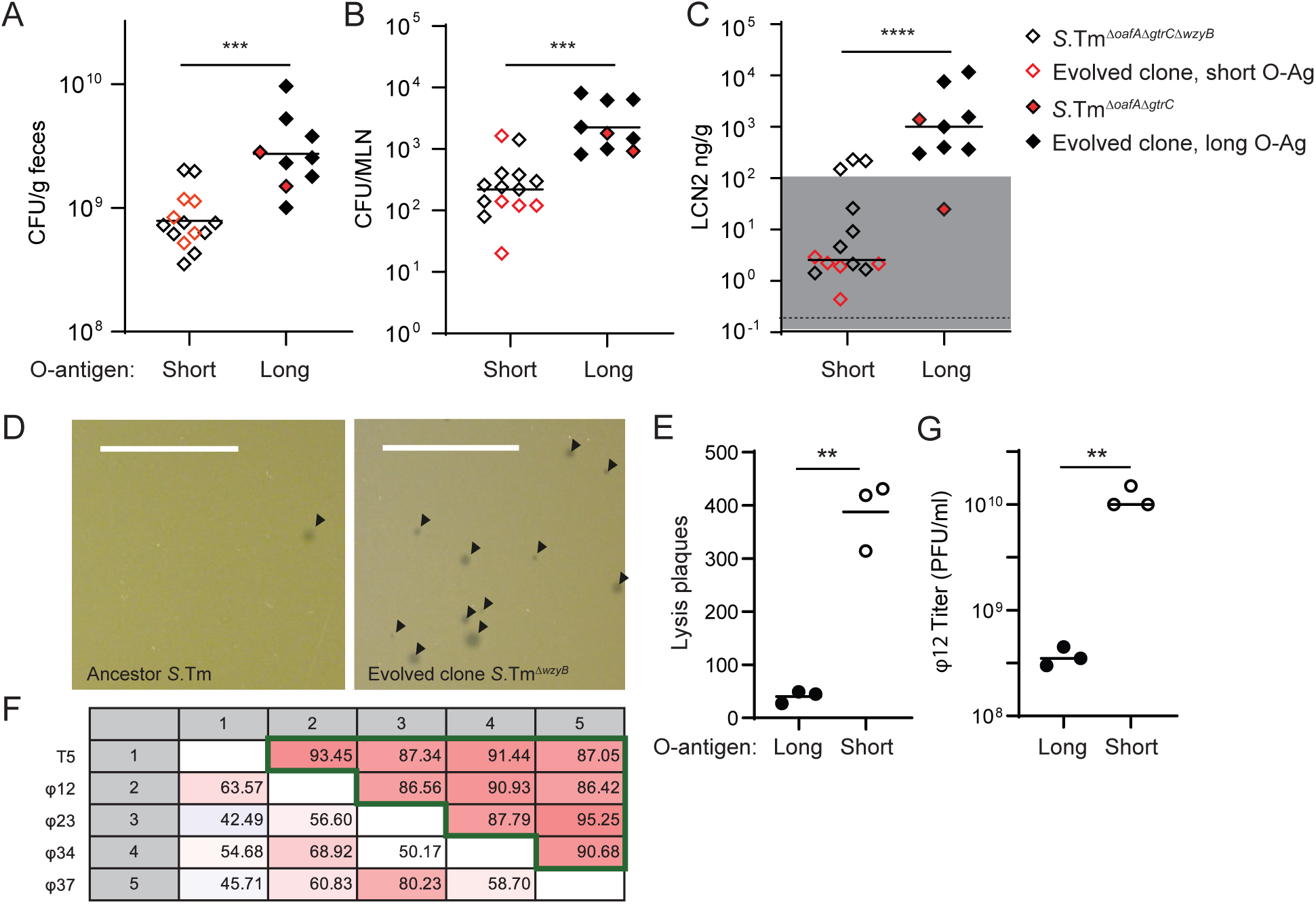
Single-repeat O-antigen mutants arising during infection of vaccinated mice have attenuated virulence, fitness and diminished resistance to phage predation. **A, B, C**, Single 24h infections in streptomycin pretreated naïve C57BL/6 mice (n=14, short O-antigen, n=9 long O-antigen). Evolved and synthetic *wzyB* mutants have reduced ability to colonize the gut (**A**, CFU/g feces, ***p=0.0002) and to spread systemically (**B**, CFU per mesenteric lymph node (MLN), ***p=0.0001). This translates into diminished propensity to trigger intestinal inflammation in comparison to isogenic wild type strains (**C**, fecal Lipocalin 2 (LCN2), ****p<0.0001). Mann-Whitney U, 2-tailed tests. **D**. Phage plaques on a lawn of ancestor *S.* Tm^wt^ (left) and evolved *S.*Tm^Δ*wzyB*^ (right) after infection with filtered wastewater; scale=1cm. **E**. Quantification of the plaques from three independent experiments (2-tailed Paired T test **p=0.0046). **F**. Pairwise comparison matrix of *de novo* assembled and aligned genomes of isolated bacteriophages (φ12, φ23, φ34, φ37) and a reference sequence from *Enterobacteriaceae* phage T5 (NC_005859). Values indicate the alignment percentage (comparisons below diagonal) between genomes and the average nucleotide identity between the aligned parts (comparisons above diagonal, green frame). This analysis shows that the four isolated bacteriophages are different but all belong to the T5 family. **G**. Quantification of phage plaques formed on infection of the ancestor *S.*Tm^wt^ (long O-antigen) and evolved *S.*Tm^Δ*wzyB*^ (short O-antigen) with the isolated phage φ12. 2-tailed Mann-Whitney U test. **p=0.0041.

These observations revealed the overlap between host IgA driven- and phage-driven *Salmonella* evolution. Both the *oafA* gene and the *gtrABC* operon are found at bacteriophage remnant loci, indicating that *S.*Tm has co-opted functions modulating sensitivity to bacteriophage attack in order to escape adaptive immunity. Of note, this example of “coincidental evolution”^31, 32^ could be also driven by and influence how *Salmonella* escape protozoa predator grazing in the gut^33^. As protozoa are specifically excluded from our SPF mouse colonies, this effect could not be investigated here.

Our data, along with previous work on O:4[5] and O:12 variation^4, 5, 9, 10^, clearly indicated direct selective pressure of the host immune system for within-host evolution/phase variation of the O-antigen. Nevertheless, IgA specificity is only one of many strong selective pressures that can be present in the intestine of a free-living animal. Previous work^20, 32–34^ indicates that inflammation, phage and predation by protozoa can all contribute, and may exhibit complex interactions. For example, inflammation induces the lytic cycle of a temperate phage: a phenomenon inhibited by IgA-mediated protection from disease^20^. Inflammation is also expected to be particularly detrimental to O-antigen-deficient strains that are poorly resistant to antimicrobial peptides and bile acids^3^ (Fig. ED9). Aggregation of *Salmonella* by IgA may also generate particles that are too large for protozoal grazing, further interacting with bacterial predation in the gut, although this hypothesis has not been experimentally tested. We hope that our work has generated a framework and a set of tools that can be applied to better understand the influence of intestinal adaptive immunity on within-host evolution of bacteria more comprehensively, and that eventually this can be translated into better control of enteric pathogens. In our case, we observed that a tailored adaptive immune response can influence the evolution of bacteriophage/bacteria interactions to the detriment of the bacteria.

We have focused on one particular *S.*Tm strain here and it remains to be seen how far this concept can be extended. Further phage-encoded modification of the O-antigen, such as the O:12-1 modification^4^ will likely be required to make robust “evolutionary traps” for *Salmonella* Typhimurium “in the wild”. Additionally, species capable of producing capsular polysaccharides that mask the O-antigen, such as *Salmonella* Typhi and many *E.coli* strains, would require additional vaccine components (typically glycoconjugates) able to induce robust anti-capsule immunity. However, we expect the principle uncovered here, i.e. understanding the rapid within-host evolution of bacterial surface structures and using this information to rationally design oligomeric vaccines, to be broadly applicable. Correspondingly, our findings are consistent with earlier reports of IgA-mediated selection of surface glycans in diverse species^14, 35^, and an earlier report that *gtrABC*-mediated O-antigen phase-variation of *Salmonella* Typhimurium ATCC 14028 confers a colonization benefit starting at day 10 post-infection (roughly the time when an IgA response would be first detected)^5^. Surface variation of teichoic acids for immune evasion can also be prophage-driven in *Staphylococcus aureus*^36^, although adaption of antibody-based techniques for gram-positive pathogens that are masters of immune evasion will likely be beyond the limits of this approach.

“Evolutionary trapping” of *Salmonella* by vaccine-induced IgA does not require any effect of IgA on the intrinsic mutation rate or phase-switching rates of *Salmonella.* Rather within-host evolution is the product of specific selective pressures (driven by IgA) on mutants and phase variants with changes in O-antigen structure, which are spontaneously generated at relatively high frequencies in the course of any intestinal infection. This genetic plasticity of large populations of microbes has always been the “Achilles heel” of antibiotic^37^, phage^38^ or CRISPR-based^39^ treatments, leading to resistance and treatment failure. In the complex ecological setting of the intestine, where bacterial populations are large and relatively fast-growing, within-host evolution can be rapid, and surprisingly predictable. Via rationally designed oral vaccines, we demonstrate that this force can be harnessed to weaken pathogenicity and to alter bacterial susceptibility to predation. We therefore propose that understanding the most common within-host evolutionary trajectories of gut pathogens holds the key to developing robust prophylactics and therapies.

## Supporting information

Numerical source data Fig. 1

numerical source data Fig. 2

numerical source data Fig. 3

numerical source data Fig. 4

Numerical Source data supplementary figures

## Acknowledgements

OH acknowledges Heiko Käßner for recording NMR spectra, Regina Engel for GLC-MS, and Katharina Jakob and Sylvia Düpow for technical support. We want to thank Magdalena Schneider, Christine Kiessling, Elisabeth Schultheiss, Rosa-Maria Vesco and Clarisse Straub for the DNA extraction, library preparations and sequencing of the bacterial isolates. MD acknowledges Delphine Cornillet and the group of Prof. Dirk Bumann for serum resistance measurements.

ES, WDH and MD acknowledge Prof. Markus Aebi, Prof. Martin Loessner, Dr. Joshua Cherry and the group of Dr. Alexander Harms for their helpful discussion and insight, as well as members of the Hardt, Slack, and Diard groups for their comments. They further acknowledge the staff of the ETH Phenomics Centre and Rodent Centre HCI for support for animal experimentation.

## Author contributions

MD, WDH and ES designed the project and wrote the paper. MD and ES designed and carried out experiments relating to vaccination and infection of mice, re-isolation of *S.*Tm clones, phenotyping of *S.*Tm clones by flow cytometry and gel electrophoresis, characterization of human monoclonal antibodies, analysis of antibody titres, and analysis of fitness of O-antigen variants of *S.*Tm *in vitro* and *in vivo*. MvdW, BHM, CL, RM contributed to experimental design / data interpretation. GZ carried out HR-MAS NMR analysis, OH carried out proton NMR analysis. MA generated the mathematical model for O:12 switching. JA carried out and analysed all AFM imaging. AR, NAB carried out phage-sensitivity assays. AE, FB, DW carried out Illumina whole-genome resequencing of re-isolated *S.*Tm isolates. EB, VL, DH, FB, KSM, SA carried out *S.*Tm challenge infections in vaccinated mice and analysed re-isolated clones. AH carried out microfluidic video microscopy of O:12 switching. PV and LF carried out methylome analysis of re-isolated *S.*Tm clones. LP, AL and BMS generated novel antibody reagents. All authors critically reviewed the manuscript.

## Materials and Methods

### Ethics statement

All animal experiments were approved by the legal authorities (licenses 223/2010, 222/2013, 193/2016, 120/2019; Kantonales Veterinäramt Zürich, Switzerland). All experiments involving animals were carried out strictly in accordance with the legal framework and ethical guidelines.

### Mice

Unless otherwise stated, all experiments used specific opportunistic pathogen-free (SPF, containing a complete microbiota free of an extended list of opportunistic pathogens) C57BL/6 mice. IgA^−/−^ ^40^, Balb/c, J_H_^-/-^ ^41^, Rag1^-/-^ ^42^ (all C57BL/6 background) and 129S1/SvImJ, mice, were re-derived into a specific pathogen-free (SPF) foster colony to normalize the microbiota and bred under full barrier conditions in individually ventilated cages in the ETH Phenomics Center (EPIC, RCHCI), ETH Zürich and were fed a standard chow diet. Low complex microbiota (LCM) mice (IgA+/- and -/-, used in Fig. ED2) are ex-germfree mice, which were colonized with a naturally diversified Altered Schaedler flora in 2007^14^ and were bred in individually ventilated cages or flexible-film isolators at this facility, and received identical diet. All mouse facilities were regulated to maintain constant temperature (22°C +/- 1°C) and humidity (30-50%), with a 12h/12h standard dark/light cycle. Male and female mice were included in all experimental groups, and the number of animals per group is indicated in each figure legend.

Vaccinations and chronic infections with attenuated *Salmonella* strains in naïve mice were started between 5 and 6 weeks of age, and males and females were randomized between groups to obtain identical ratios wherever possible. Challenge infections with virulent *Salmonella* were carried out between 9 and 12 weeks of age. As strong phenotypes were expected, we adhered to standard practice of analysing at least 5 mice per group. Researchers were not blinded to group allocation.

### Strains and plasmids

All strains and plasmids used in this study are listed **Table S1**.

For cultivation of bacteria, we used lysogeny broth (LB) containing appropriate antibiotics (i.e., 50 µg/ml streptomycin (AppliChem); 6 µg/ml chloramphenicol (AppliChem); 50 µg/ml kanamycin (AppliChem); 100 µg/ml ampicillin (AppliChem)). Dilutions were prepared in Phosphate Buffer Saline (PBS, Difco). In-frame deletion mutants (e.g. *gtrC::cat*) were performed by *λ red* recombination as described in^43^. When needed, antibiotic resistance cassettes were removed using the temperature-inducible FLP recombinase encoded on pCP20^43^. Mutations coupled with antibiotic resistance cassettes were transferred into the relevant genetic background by generalized transduction with bacteriophage P22 HT105/1 *int-201*^44^. Primers used for genetic manipulations and verifications of the constructions are listed **Table S2.** Deletions of *gtrA* and *gtrC* originated from in-frame deletions made in *S*.Tm 14028S, kind gifts from Prof. Michael McClelland (University of California, Irvine), and were transduced into the SB300 genetic background.

The *gtrABC* operon (STM0557-0559) was cloned into the pSC101 derivative plasmid pM965^45^ for constitutive expression. The operon *gtrABC* was amplified from the chromosome of SB300 using the Phusion Polymerase (ThermoFisher Scientific) and primers listed **Table S2**. The PCR product and pM965 were digested with PstI-HF and EcoRV-HF (NEB) before kit purification (SV Gel and PCR Clean up System, Promega) and ligation in presence of T4 ligase (NEB) following manufacturer recommendations. The ligation product was transferred by electro-transformation in competent SB300 cells.

### Targeted sequencing

Targeted re-sequencing by the Sanger method (Microsynth AG) was performed on kit purified PCR products (Promega) from chromosomal DNA or expression vector templates using pre-mixed sequencing primers listed **Table S2**.

### Whole-genome re-sequencing of O:12^Bimodal^ isolates

The genomes of S.Tm and evolved derivatives were fully sequenced by the Miseq system (2×300bp reads, Illumina, San Diego, CA) operated at the Functional Genomic Center in Zürich. The sequence of *S*.Tm SL1344 (NC_016810.1) was used as reference. Quality check, reads trimming, alignments, SNPs and indels calling were performed using the bioinformatics software CLC Workbench (Qiagen).

### Whole-genome sequencing of *S*.Tm isolates from “Evolutionary Trap” vaccinated mice and variant calling

Nextera XT libraries were prepared for each of the samples. The barcoded libraries were pooled into equimolar concentrations following manufacturer’s guidelines (Illumina, San Diego, CA) using the Mid-Output Kit for paired-end sequencing (2×150 bp) on an Illumina NextSeq500 sequencing platform. Raw data (mean virtual coverage 361x) was demultiplexed and subsequently clipped of adapters using Trimmomatic v0.38 with default parameters^46^. Quality control passing read-pairs were aligned against reference genome/plasmids (Accession numbers: NC_016810.1, NC_017718.1, NC_017719.1, NC_017720.1) with bwa v0.7.17^47^. Genomic variant were called using Pilon v1.23^48^. with the following parameters: (i) minimum coverage 10x; (ii) minimum quality score = 20; (iii) minimum read mapping quality = 10. SnpEff v4.3 was used to annotate variants according to NCBI and predict their effect on genes^49^.

### PA-STm vaccinations

Peracetic acid killed vaccines were produced as previously described^28^. Briefly, bacteria were grown overnight to late stationary phase, harvested by centrifugation and re-suspended to a density of 10^9^-10^10^ per ml in sterile PBS. Peracetic acid (Sigma-Aldrich) was added to a final concentration of 0.4% v/v. The suspension was mixed thoroughly and incubated for 60 min at room temperature. Bacteria were washed once in 40 ml of sterile 10x PBS and subsequently three times in 50 ml sterile 1x PBS. The final pellet was re-suspended to yield a density of 10^11^ particles per ml in sterile PBS (determined by OD600) and stored at 4°C for up to three weeks. As a quality control, each batch of vaccine was tested before use by inoculating 100 µl of the killed vaccine (one vaccine dose) into 300 ml LB and incubating over night at 37 °C with aeration. Vaccine lots were released for use only when a negative enrichment culture had been confirmed. For all vaccination, 10^10^ particles, suspended in 100µl PBS were delivered by oral gavage, once weekly for 4 weeks. Where multiple strains were combined, the total number of vaccine particles remained constant, and was roughly equally divided between the constituent strains. Unless otherwise stated, PA-STm vaccinated mice were challenged orally on d28 after the first vaccination.

### Adoptive transfer of recombinant mSTA121 IgA

A recombinant monoclonal dimeric murine IgA specific for the O:12-0 epitope (described in ^15^) was buffer-exchanged into sterile PBS. 1 mg of antibody was injected intravenously into mice 30 min prior to infection and again 12 h post-infection, to maintain sufficient dimeric IgA for export into the gut by PIgR.

### Chronic infection with live-attenuated vaccine strains of non-typhoidal *Salmonella*

6-week-old mice were orally pretreated 24 h before infection with 25 mg streptomycin. Live-attenuated strains *(sseD::aphT*, *ΔgtrC ΔaroA* and *ΔoafA ΔgtrC ΔaroA*, **Table S1**,^50^) were cultivated overnight separately in LB containing streptomycin. Subcultures were prepared before infections by diluting overnight cultures 1:20 in fresh LB without antibiotics and incubation for 4 h at 37°C. The cells were washed in PBS, diluted, and 50 µl of resuspended pellets were used to infect mice *per os* (5×10^7^ CFU).

Feces were sampled at day 1, 9 and 42 post-infection, homogenized in 1 ml PBS by bead beating (3mm steel ball, 25 Hz for 1 minute in a TissueLyser (Qiagen)), and *S*.Tm strains were enumerated by selective plating on MacConkey agar supplemented with streptomycin. Samples for lipocalin-2 measurements were kept homogenized in PBS at −20 °C. Enrichment cultures for analysis of O-antigen composition were carried out by inoculating 2 µl of fecal slurry into 5ml of fresh LB media and cultivating overnight at 37 °C.

### Non-typhoidal *Salmonella* challenge infections

Infections were carried out as previously described ^23^. In order to allow reproducible gut colonization, 8-12 week-old SPF mice, naïve or PA-STm vaccinated, were orally pretreated 24 h before infection with 25 mg streptomycin or 20 mg of ampicillin. Strains were cultivated overnight separately in LB containing the appropriate antibiotics. Subcultures were prepared before infections by diluting overnight cultures 1:20 in fresh LB without antibiotics and incubation for 4 h at 37°C. The cells were washed in PBS, diluted, and 50 µl of resuspended pellets were used to infect mice *per os* (5×10^5^ CFU). Competitions were performed by inoculating 1:1 mixtures of each competitor strain. Feces were sampled daily, homogenized in 1 ml PBS by bead beating (3 mm steel ball, 25 Hz for 1 min in a TissueLyser (Qiagen)), and *S*.Tm strains were enumerated by selective plating on MacConkey agar supplemented with the relevant antibiotics. Fecal samples for lipocalin-2 measurements were kept homogenized in PBS at −20°C. At endpoint, intestinal lavages were harvested by flushing the ileum content with 2 ml of PBS using a cannula. The mesenteric lymph nodes, were collected, homogenized in PBS Tergitol 0.05% v/v at 25 Hz for 2 min, and bacteria were enumerated by selective plating.

Competitive indexes were calculated as the ratio of population sizes of each genotype, enumerated by selective plating of the two different strains on kanamycin- and chloramphenicol-containing agar, at a given time point, normalized for the ratio determined by selective plating in the inoculum (which was always between 0.5 and 2).

### Non-typhoidal *Salmonella* transmission

Donor mice were vaccinated with PA-*S.*Tm^Δ*oafA* Δ*gtrC*^ once per week for 5 weeks, streptomycin pretreated (25 mg streptomycin *per os*), and gavaged 24 h later with 10^5^ CFU of a 1:1 mixture of *S.* Tm^Δ*oafA*Δ*gtrCwzyB::cat*^ (Cm^R^) and *S*. Tm^Δ*oafA*Δ*gtrC* Kan^ (Kan^R^). On day 4 post infection, the donor mice were euthanized, organs were harvested, and fecal pellets were collected, weighed and homogenized in 1 ml of PBS. The re-suspended feces (centrifuged for 10 s to discard large debris) were immediately used to gavage (as a 50 µl volume containing the bacteria from on fecal pellet) recipient naïve mice (pretreated with 25 mg streptomycin 24 hours before infection). Recipient mice were euthanized and organs were collected on day 2 post transmission. In both donor and recipient mice, fecal pellets were collected daily and selective plating was used to enumerate *Salmonella* and determine the relative proportions (and consequently the competitive index) of both competing bacterial strains.

### Quantification of fecal Lipocalin2

Fecal pellets collected at the indicated time-points were homogenized in PBS by bead-beating at 25 Hz, 1min. Large particles were sedimented by centrifugation at 300 *g*, 1 min. The resulting supernatant was then analysed in serial dilution using the mouse Lipocalin2 ELISA duoset (R&D) according to the manufacturer’s instructions.

### Analysis of specific antibody titres by bacterial flow cytometry

Specific antibody titres in mouse intestinal washes were measured by flow cytometry as described^15, 51^. Briefly, intestinal washes were collected by flushing the small intestine with 2 ml PBS, centrifuged at 16000 *g* for 30 min to clear all bacterial-sized particles. Aliquots of the supernatants were stored at −20°C until analysis. Bacterial targets (antigen against which antibodies are to be titred) were grown to late stationary phase or the required OD in 0.2µm-filtered LB, then gently pelleted for 2 min at 7000 *g*. The pellet was washed with 0.2µm-filtered 1% BSA/PBS before re-suspending at a density of approximately 10^7^ bacteria per ml. After thawing, intestinal washes were centrifuged again at 16000 *g* for 10 min to clear. Supernatants were used to perform serial dilutions. 25 μl of the dilutions were incubated with 25 μl bacterial suspension at 4°C for 1 h. Bacteria were washed twice with 200 μl 1% BSA/PBS by centrifugation at 7000g for 15 min, before resuspending in 25 μl of 0.2µm-filtered 1% BSA/PBS containing monoclonal FITC-anti-mouse IgA (BD Pharmingen, 10 µg/ml) or Brilliant violet 421-anti-IgA (BD Pharmingen, 10µg/ml). After 1 h of incubation, bacteria were washed once with 1% BSA/PBS as above and resuspended in 300 μl 1% BSA/PBS for acquisition on LSRII or Beckman Coulter Cytoflex S using FSC and SSC parameters to threshold acquisition in logarithmic mode. Data were analysed using FloJo (Treestar). After gating on bacterial particles, log-median fluorescence intensities (MFI) were plotted against lavage dilution factor for each sample and 4-parameter logistic curves were fitted using Prism (Graphpad, USA). Titers were calculated from these curves as the dilution factor giving an above-background signal (typically IgA coating MFI=1000 – e.g. Fig. S7 and S8).

### Dirty-plate ELISA analysis of intestinal lavage IgA titres specific for *S.*Tm

Bacterial targets (antigen against which antibodies are to be titred) were grown to late stationary phase in 0.2µm-filtered LB, then gently pelleted for 2 min at 7000 *g*. The pellet was washed with 0.2µm-filtered 1% BSA/PBS before re-suspending at a density of approximately 10^9^ bacteria per ml in sterile PBS. 50µl of this bacterial suspension was added to each well of a Nunc Immunosorb ELISA plate and was incubated overnight at 4°C in a humidified chamber. The ELISA plates were then washed 3 times with PBS/0.5% Tween-20 and blocked with 200µl per well of 2% BSA in PBS for 3h. After thawing, intestinal washes were centrifuged again at 16000 *g* for 10 min to clear. Supernatants were used to perform serial dilutions. 50 μl of the dilutions were added to each well and the plates were incubated at 4°C overnight in a humidified chamber. The next morning, the plates were washed 5 times with PBS/0.5% Tween-20 and 50µl of HRP-anti-mouse-IgA (Sigma-Aldrich, 1:1000) was added to each well. This was incubated for 1h at room temperature before washing again 5 times and developing the plates with 100µl per well of ABTS ELISA substrate. Absorbance at 405nm was read using a Tecan Infinite pro 200. A_405_ readings were plotted against lavage dilution factor for each sample and 4-parameter logistic curves were fitted using Prism (Graphpad, USA). Titers were calculated from these curves as the dilution factor giving an above-background signal (A_405_=0.2 – e.g. Fig. S7 and S8).

### Flow cytometry for analysis of O:5, O:4 and O:12-0 epitope abundance on *Salmonella* in cecal content, enrichment cultures and clonal cultures

1 µl of overnight cultures made in 0.2µm-filtered LB, or 1µl of fresh feces or cecal content suspension (as above) was stained with 0.2µm-filtered solutions of STA5 (human recombinant monoclonal IgG2 anti-O:12-0, 6µg/ml ^15^), Rabbit anti-*Salmonella* O:5 (Difco, 1:200) or Rabbit anti-*Salmonella* O:4 (Difco, 1:5). After incubation at 4°C for 30 min, bacteria were washed twice by centrifugation at 7000g and resuspension in PBS/1% BSA. Bacteria were then resuspended in 0.2µm-filtered solutions of appropriate secondary reagents (Alexa 647-anti-human IgG, Jackson Immunoresearch 1:200, Brilliant Violet 421-anti-Rabbit IgG, Biolegend 1:200). This was incubated for 10-60 min before cells were washed as above and resuspended for acquisition on a BD LSRII or Beckman Coulter Cytoflex S. A media-only sample was run on identical settings to ensure that the flow cytometer was sufficiently clean to identify bacteria without the need for DNA dyes. Median fluorescence intensity corresponding to O:12-0 or O:5 staining was calculated using FlowJo (Treestar, USA). Gates used to calculate the % of “ON” and “OFF” cells were set by gating on samples with known O:5/O:4 (*oafA*-deletion) and O:12-0 (*gtrC-*deletion) versus O:12-2 (*pgtrABC*) phenotypes (Fig. S2 and 3).

### Live-cell immunofluorescence

200 uL of an overnight culture was centrifuged and resuspended in 200 μL PBS containing 1 μg recombinant murine IgA clone STA121-AlexaFluor568. The cells and antibodies were co-incubated for 20 min at room temperature in the dark and then washed twice in 1 mL Lysogeny broth (LB). Antibody-labeled cells were pipetted into an in-house fabricated microfluidic device^52^. Cells in the microfluidic device were continuously fed *S*.Tm-conditioned LB^52^ containing STA121-AlexaFluor568 (1 μg/mL). Media was flowed through the device at a flow rate of 0.2 mL/h using syringe pumps (NE-300, NewEra PumpSystems). Cells in the microfluidic device were imaged on an automated Olympus IX81 microscope enclosed in an incubation chamber heated to 37°C. At least 10 unique positions were monitored in parallel per experiment. Phase contrast and fluorescence images were acquired every 3 min. Images were deconvoluted in MatLab^53^. Videos are compressed to 7 fps, i.e. 1 s = 21 mins.

## HR-MAS NMR

*S*. Typhimurium cells were grown overnight (∼18 h) to late stationary phase. The equivalent of 11–15 OD_600_ was pelleted by centrifugation for 10 min 4 °C and 3750 *g*. The pellet was resuspended in 10% NaN_3_ in potassium phosphate buffer (PPB; 10 mM pH 7.4) in D_2_O and incubated at room temperature for at least 90 min. The cells were then washed twice with PPB and resuspended in PPB to a final concentration of 0.2 OD_600_/μl in PPB containing acetone (final concentration 0.1% (v/v) as internal reference). The samples were kept on ice until the NMR measurements were performed -i.e. for between 1 and 8 h. The HR-MAS NMR spectra were recorded in two batches, as follows: *S*.Tm^WT^, *S*.Tm^Δ*wbaP*^, *S*.Tm^Evolved_1^, *S*.Tm^Evolved_2^ were measured on 16.12.2016, *S*.Tm^Δ*oafA*^ was measured on 26.7.2017.

NMR experiments on intact cells were carried out on a Bruker Biospin AVANCE III spectrometer operating at 600 MHz ^1^H Larmor frequency using a 4 mm HR-MAS Bruker probe with 50 µl restricted-volume rotors. Spectra were collected at a temperature of 27 °C and a spinning frequency of 3 kHz except for the sample of *S*.Tm^Δ*oafA*^ (25°C, 2 kHz). The ^1^H experiments were performed with a 24 ms Carr– Purcell–Meiboom–Gill (CPMG) pulse-sequence with rotor synchronous refocusing pulses every two rotor periods before acquisition of the last echo signal to remove broad lines due to solid-like material^54^. The 90° pulse was set to 6.5 μs, the acquisition time was 1.36 s, the spectral width to 20 ppm. The signal of HDO was attenuated using water pre-saturation for 2 s. 400 scans were recorded in a total experimental time of about 30 minutes.

### O-Antigen purification and ^1^H-NMR

The LPS was isolated applying the hot phenol-water method^55^, followed by dialysis against distilled water until the phenol scent was gone. Then samples were treated with DNase (1mg/100 mg LPS) plus RNase (2 mg/100 mg LPS) at 37°C for 2 h, followed by Proteinase K treatment (1 mg/100 mg LPS) at 60°C for 1 h [all enzymes from Serva, Germany]. Subsequently, samples were dialyzed again for 2 more days, then freeze dried. Such LPS samples were then hydrolyzed with 1% aqueous acetic acid (100°C, 90 min) and ultra-centrifuged for 16 h at 4°C and 150,000 *g*. Resulting supernatants (the O-antigens) were dissolved in water and freeze-dried. For further purification, the crude O-antigen samples were chromatographed on TSK HW-40 eluted with pyridine/acetic acid/water (10/4/1000, by vol.), then lyophilized. On these samples, 1D and 2 D (COSY, TOCSY, HSQC, HMBC) ^1^H- and ^13^C-NMR spectra were recorded with a Bruker DRX Avance 700 MHz spectrometer (^1^H: 700.75 MHz; ^13^C: 176.2 MHz) as described^56^.

### Atomic force microscopy

The indicated *S*.Tm strains were grown to late-log phase, pelleted, washed once with distilled water to remove salt. A 20 µl of bacterial solution was deposited onto freshly cleaved mica, adsorbed for 1 min and dried under a clean airstream. The surface of bacteria was probed using a Dimension FastScan Bio microscope (Bruker) with Bruker AFM cantilevers in tapping mode under ambient conditions. The microscope was covered with an acoustic hood to minimized vibrational noise. AFM images were analyzed using the Nanoscope Analysis 1.5 software.

### Methylation analysis of *S*.Tm clones

For REC-Seq (restriction enzyme cleavage–sequencing) we followed the same procedure described by Ardissone et al, 2016^57^. In brief, 1 µg of genomic DNA from each S.Tm was cleaved with MboI, a blocked (5’biotinylated) specific adaptor was ligated to the ends and the ligated fragments were then sheared to an average size of 150-400 bp (Fasteris SA, Geneva, CH). Illumina adaptors were then ligated to the sheared ends followed by deep-sequencing using a HiSeq Illumina sequencer, the 50 bp single end reads were quality controlled with FastQC v0.9 (http://www.bioinformatics.babraham.ac.uk/projects/fastqc/). To remove contaminating sequences, the reads were split according to the MboI consensus motif (5’-^GATC-3’) considered as a barcode sequence using fastx_toolkit v0.0.13.2 (http://hannonlab.cshl.edu/fastx_toolkit/) (fastx_barcode_splitter.pl --bcfile barcodelist.txt --bol --exact). A large part of the reads (60%) were rejected and 40% kept for remapping to the reference genomes with bwa mem^47^ v0.7.15 and samtools^58^ v0.1.19 to generate a sorted bam file. The bam file was further filtered to remove low mapping quality reads (keeping AS >= 45) and split by orientation (alignmentFlag 0 or 16) with bamtools^59^ v2.4.1. The reads were counted at 5’ positions using Bedtools^60^ v2.26.0 (bedtools genomecov -d -5). Both orientation count files were combined into a bed file at each identified 5’-GATC-3’ motif using PERL script (perl v5.24). The MboI positions in the bed file were associated with the closest gene using bedtools closest^60^ v2.26.0 and the gff3 file of the reference genomes^61^. The final bed file was converted to an MS Excel sheet. The counts were loaded in RStudio v1.1.442^62^ with R v3.4.4^63^ and analysed with the DESeq2 v1.18.1 package^64^ comparing the reference strain with the 3 evolved strains considered as replicates. The counts are analysed by genome position rather than by gene. The positions are considered significantly differentially methylated upon an adjusted p-value < 0.05. Of the 2607 GATC positions, only 4 were found significantly differentially methylated and they are all located in the promoter of the *gtrABC* operon.

The first step in the reads filtering was to remove contaminant reads missing the GATC consensus motif (MboI) at the beginning of the sequence. These contaminant reads are due to random fragmentation of the genomic DNA and not to cuts of the MboI restriction enzyme. Using fastx_barcode_splitter.pl v0.0.13.2 about 60% of the reads were rejected because they did not start with GATC. The rest (40%) was analyzed further. Random DNA shearing and blunt-ended ligation of adaptors, combined with sequencing noise at the beginning of reads likely generates this high fraction of reads missing at GTAC sequence.

### *gtrABC* expression analysis by blue/white screening and flow cytometry

About 200 colonies of *S*.Tm*^gtrABC-lacZ^* (strain background 4/74, ^4^) were grown from an overnight culture on LB agar supplemented with X-gal (0.2 mg/ml, Sigma) in order to select for *gtrABC* ON (blue) and OFF clones (white). These colonies were then picked to start pure overnight cultures. These cultures were diluted and plated on fresh LB agar X-gal plate in order to enumerate the proportion of *gtrABC* ON and OFF siblings. The proportion of O:12/O:12-2 cells was analyzed by flow cytometry.

### *In vitro* growth and competitions to determine *wzyB*-associated fitness costs

Single or 1:1 mixed LB subcultures were diluted 1000 times in 200 µl of media distributed in 96 well black side microplates (Costar). Where appropriate, wild type *S.*Tm carried a plasmid for constitutive expression of GFP. To measure growth and competitions in stressful conditions that specifically destabilize the outer membrane of *S*.Tm, a mixture of Tris and EDTA (Sigma) was diluted to final concentration (4 mM Tris, 0.4 mM EDTA) in LB; Sodium cholate (Sigma) and Sodium Dodecyl Sulfate (SDS) (Sigma) were used at 2% and 0.05% final concentration respectively. The lid-closed microplates were incubated at 37°C with fast and continuous shaking in a microplate reader (Synergy H4, BioTek Instruments). The optical density was measured at 600 nm and the green fluorescence using 491 nm excitation and 512 nm emission filter wavelengths every 10 minutes for 18 h. Growth in presence of SDS causes aggregation when cell density reaches OD=0.3-0.4, therefore, it is only possible to compare the growth curves for about 250 minutes. The outcome of competitions was determined by calculating mean OD and fluorescence intensity measured during the last 100 min of incubation. OD and fluorescence values were corrected for the baseline value measured at time 0.

### Serum resistance

Overnight LB cultures were washed three times in PBS, OD adjusted to 0.5 and incubated with anonymized pooled human serum obtained from Unispital Basel (3 vol of culture for 1 vol of serum) at 37°C for 1 h. Heat inactivated (56°C, 30 min) serum was used as control treatment. Surviving bacteria were enumerated by plating on non-selective LB agar plates. For this, dilutions were prepared in PBS immediately after incubation.

### Bacteriophage sensitivity tests

5 ml sewage water (sewage plant inflow treated with 1 % *v/v* chloroform; Basel Stadt, Switzerland) were mixed with 500 µl of dense bacterial culture (ancestor wild type *S*. Tm; evolved short O-antigen *wzyB* mutant AE860.3, *S*.Tm *^ΔgtrC oafA::cat^*, *S*.Tm *^ΔgtrC ΔoafA wzyB::cat^*), incubated for 15 minutes at 37 °C. The mixtures were added to 15 ml LB containing 10 mM CaCl_2_, 10 mM MgSO_4_ and 0.7 % *w/v* agar, and immediately poured onto LB agar plates with the appropriate antibiotics.

Sensitivity to isolated phage φ12 was quantified by calculating phage titres obtained after overnight cultures of evolved short O-antigen *wzyB* mutant AE860.3 or ancestor wild type *S*. Tm in presence of the isolated bacteriophage (MOI=10).

### Isolation of bacteriophages and resistant clones

Plaques with different morphologies appearing on *S*.Tm*^ΔgtrC ΔoafA wzyB::cat^* plates were streaked on overlay plates containing *S*.Tm*^ΔgtrC ΔoafA wzyB::cat^*. The resulting plaques were used to inoculate 200 µl of a *S*.Tm*^ΔgtrC ΔoafA wzyB::cat^* culture at OD_600_=0.3 in a 96-well plate and optical density was measured every 10 minutes at 37 °C with shaking in a Synergy 2 plate-reader. Well contents after 18 hours of growth were streaked onto LB-Cm plates to isolate bacterial colonies from the regrowing population. Resistance to phage was confirmed by testing for absence of plaque formation in presence of the corresponding phage.

The rest of the well contents were cleared by centrifugation and filtered (0.45 μm) for phage purification. The cleared supernatants were used to inoculate 20 ml of a *S*.Tm*^ΔgtrC ΔoafA wzyB::cat^* culture at OD_600_=0.3 and subsequently grown at 37 °C for 5 hours. Cell debris was removed by centrifugation, the supernatants cleared by 0.45 μm filtration and stored at 4 °C.

### Phage genome sequencing and analysis

Phage DNA was isolated using the Phage DNA Isolation Kit from Norgen Biotek and sequenced at MiGS, Pittsburgh, Pennsylvania, USA. For this, Nextera libraries were prepared for each sample and sequenced on an Illumina NextSeq 550 sequencing platform to generate paired end reads.

De novo genome assembly was performed using the De Novo Assembly Algorithm of CLC Genomics Workbench and the resulting high coverage contigs were aligned using the Whole Genome Alignment Plug-In to calculate neighbor-joining trees and corresponding pairwise comparison tables.

Assembly of the phage genomes resulted in a single contig of 108,227 bp and 114,055 bp for φ12 and φ23, respectively (4,928 and 4,495-fold coverage). For φ34 four separate contigs with more than 3000-fold coverage were identified (81,319, 12,250, 10,937, 5,594 bp), giving a total genome size of more than 100,100 bp, while for φ37 three contigs with more than 1600-fold coverage (95,133, 14,559, 4,197 bp) gave a total genome size of at least 113,889 bp.

For comparison, enterobacteria phage T5 has a double-stranded linear DNA genome of 121,750 bp.

### Modeling antigen switching between O12 and O12-2

The aim of this modeling approach is to test whether a constant switching rate between an O12 and an O12-2 antigen expression state can explain the experimentally observed bimodal populations.

To this end, we formulated a deterministic model of population dynamics of the two phenotypic states as:

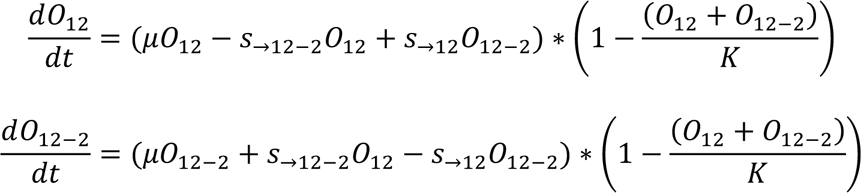

where *O*_12_ and *O*_12-2_ denote the population sizes of the respective antigen variants, *μ* denotes the growth rate, which is assumed to be identical for the two variants, *K* the carrying capacity, and *s*_→12-2_ and *s*_→12_ the respective switching rates from *O*_12_ to *O*_12-2_ and from *O*_12-2_ to *O*_12_. Growth, as well as the antigen switching rates, are scaled with population size in a logistic way, so that all processes come to a halt when carrying capacity is reached.

We use the model to predict the composition of a population after growth in LB overnight, and therefore set the specific growth rate to *μ* = 2.05ℎ^-1^, which corresponds to a doubling time of roughly 20min. The carrying capacity is set to *K* = 10^9^ *cells* . We ran parameter scans for the switching rates *s*_→12_ and *s*_→12-2_, with population compositions that start either with 100% or 0% *O*_12_, and measure the composition of the population after 16h of growth (**Fig. S11C**). The initial population size is set to 10^4^ *cells*

Experimentally, we observe that when starting a culture with an *O*_12_ colony, after overnight growth the culture is composed of around 90% *O*_12_ and 10% *O*_12-2_ cells, whereas starting the culture with *O*_12-2_ cells yields around 50% *O*_12_ and 50% *O*_12-2_ cells after overnight growth (**Fig. S11B**). To explain this observation without a change in switching rates, we would need a combination of values in *s*_→12_ and *s*_→12-2_ that yield the correct population composition for both scenarios. In **Fig. S11D**, we plot the values of *s*_→12_ and *s*_→12-2_ that yield values of 10% *O*_12-2_ (starting with 0% *O*_12-2_, green dots) and 50% *O*_12-2_ (starting with 100% *O*_12-2_, orange dots). The point clusters intersect at *s*_→12_ = 0.144ℎ^-1^ and *s*_→12-2_ = 0.037ℎ^-1^ (as determined by a local linear regression at the intersection point).

We then used the thus determined switching rates to produce a population growth curve in a in a deterministic simulation, using the above equations for a cultures starting with 100% *O*_12-2_, (**Fig. S11E**, Left-hand graph) and for a culture starting with 0% *O*_12-2_ (**Fig. S11E**, right-hand graph).

These switching rates are consistent with published values ^4^. Our results show that the observed phenotype distributions can be explained without a change in the rate of switching between the phenotypes.

## Data availability

All Plotted data and associated raw numerical data and calculations for figure 1-4, extended data fig. 1-10 and supplementary figures 1-10 is provided in source data tables (one per figure, titled accordingly). Uncropped images are provided as supplementary files.

All raw flow cytometry data, ordered by figure, is publically available via the ETH research collection doi: 10.3929/ethz-b-000477737

All Illumina sequencing data data is publically available at NCBI BioProject Accession: PRJNA720270

## Code availability

R code used to generate the figures shown in extended data figure 5 can be freely downloaded from https://github.com/marnoldini/evotrap

---

**Fig. ED1:**
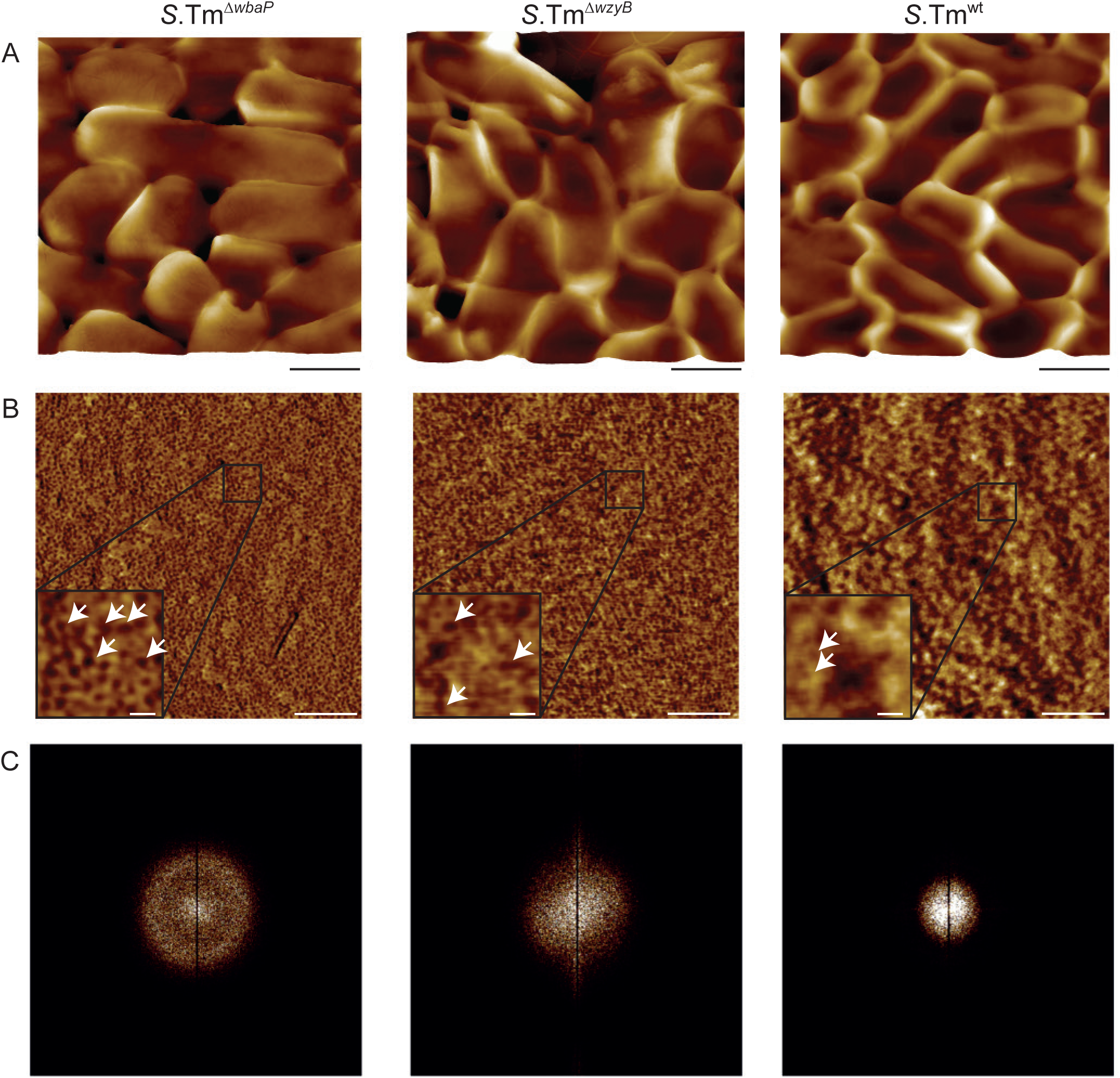
Surface phenotype of *S.*Tm mutants. **A-C**. Atomic force microscopy phase images of *S.*Tm^wt^, *S.*Tm^Δ*wzyB*^ (single-repeat O-antigen), and *S.*Tm^Δ*wbaP*^ (rough mutant -no O-antigen) at low magnification (A, uncropped image, scale bar = 1µm) and high magnification (B and C, scale bar main image = 150nm, scale bar inset = 15nm). Invaginations in the surface of *S.*Tm^Δ*wbaP*^ (dark colour, B) show a geometry and size consistent with outer membrane pores^65^. These are already less clearly visible on the surface of *S.*Tm^Δ*wzyB*^ with a single-repeat O-antigen, and become very difficult to discern in *S.*Tm^wt^. One representative image of 3 for each genotype is shown. While arrows point to features with consistent size and abundance to be exposed outer membrane porins. **C**. Fast-Fourier transform of images shown in “B” demonstrating clear regularity on the surface of *S.*Tm^Δ*wbaP*^, which is progressively lost when short and long O-antigen is present.

**Fig. ED2:**
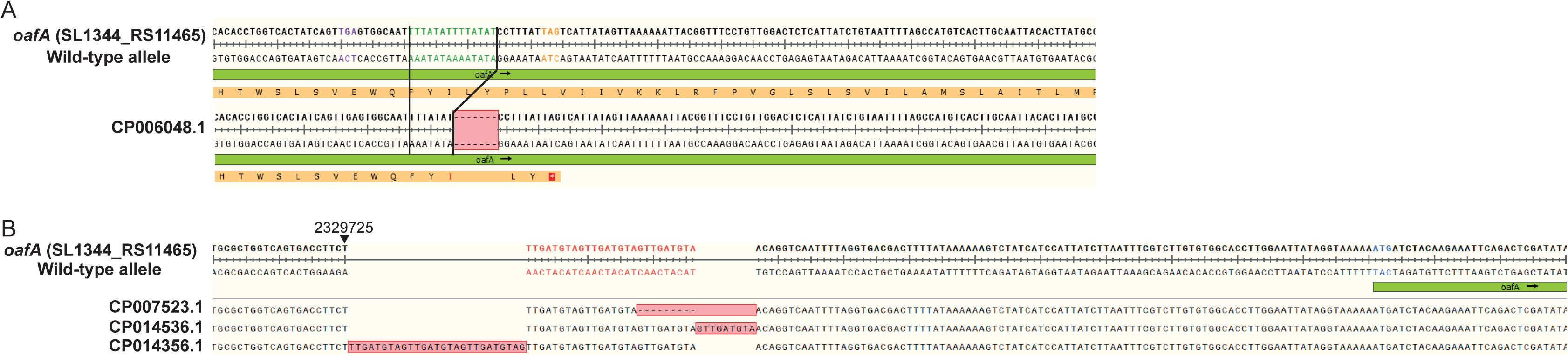
Mutations detected in the *oafA* gene sequence among several strains of *S.*Tm. **A**. Aligned fractions of the *oafA* ORF from a natural isolate (from chicken) presenting the same 7 bp deletion detected in mutants of *S*.Tm SL1344 emerging in vaccinated mice. *S*.Tm SL1344 was used a reference^66^. **B**. Aligned *oafA* promoter sequences from three natural isolates of human origin (stool or cerebrospinal fluid^67^) showing variations in the number of 9 bp direct repeats.

**Fig. ED3:**
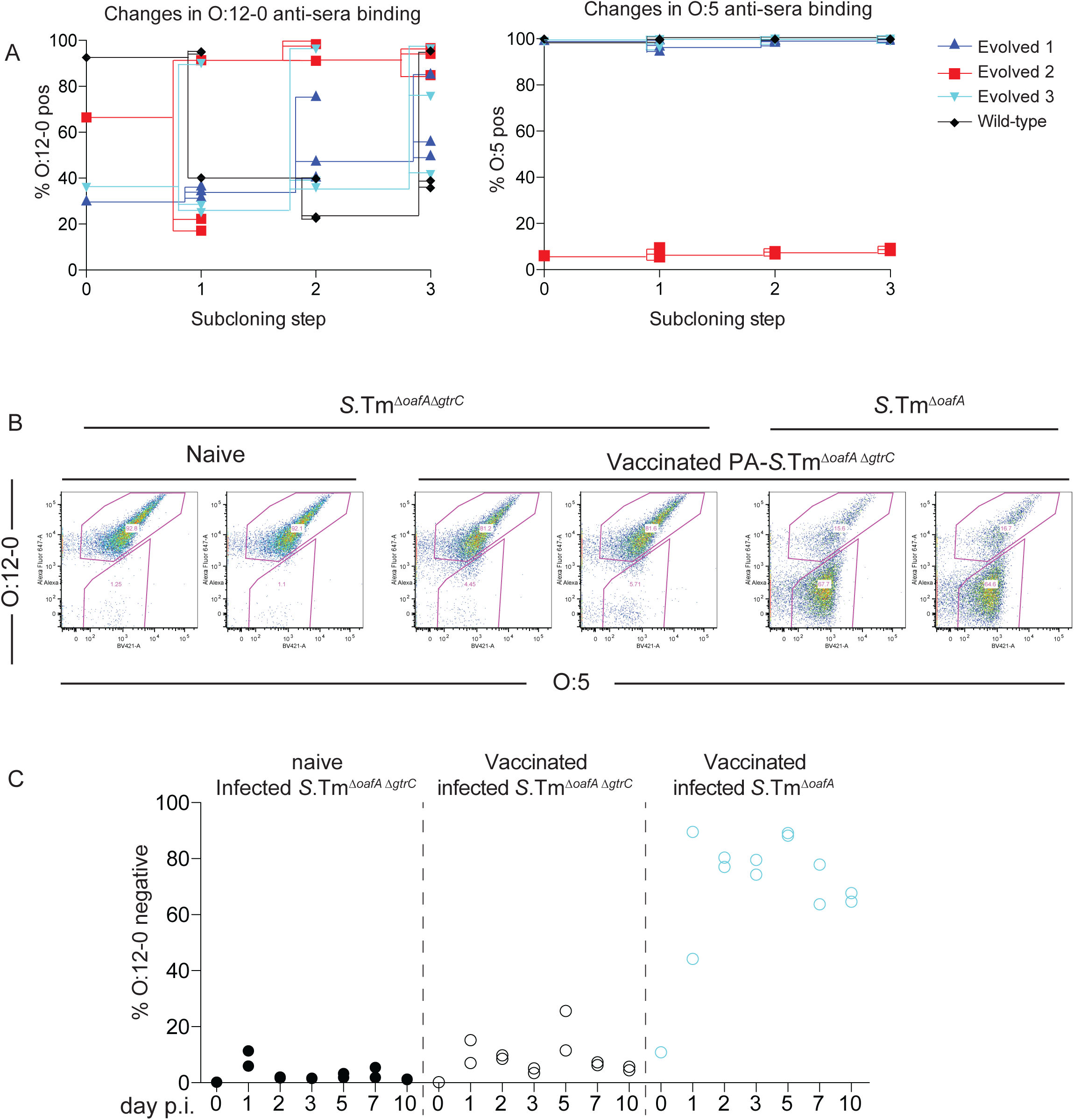
Loss of the O:12-0 epitope is a reversible phenotype. **A.** Wild type and evolved *S.*Tm clones were picked from LB plates, cultured overnight, phenotypically characterized by O:12-0 (left panel) and O:5 staining (right panel), plated and re-picked. This process was repeated over 3 cycles with lines showing the descendants of each clone. **B and C.** Wild type 129S1/SvImJ mice were mock-vaccinated or were vaccinated with PA-*S.*Tm^Δ*oafA* Δ*gtrC*^ as in Fig. 1. On d28, all mice were pre-treated with streptomycin, and infected with the indicated strain. **B**. Feces recovered at day 10 post-infection, was enriched overnight by culture in streptomycin, and stained for O:12-0 (human monoclonal STA5). Fraction O:12-0-low *S*.Tm was determined by flow cytometry. Percentage of *S.*Tm that are O:12-0-negative was quantified over 10 days and is plotted in panel **C**. Vaccination selects for *S.*Tm that have lost the O:12-0 epitope, only if the *gtrC* gene is intact.

**Fig. ED4:**
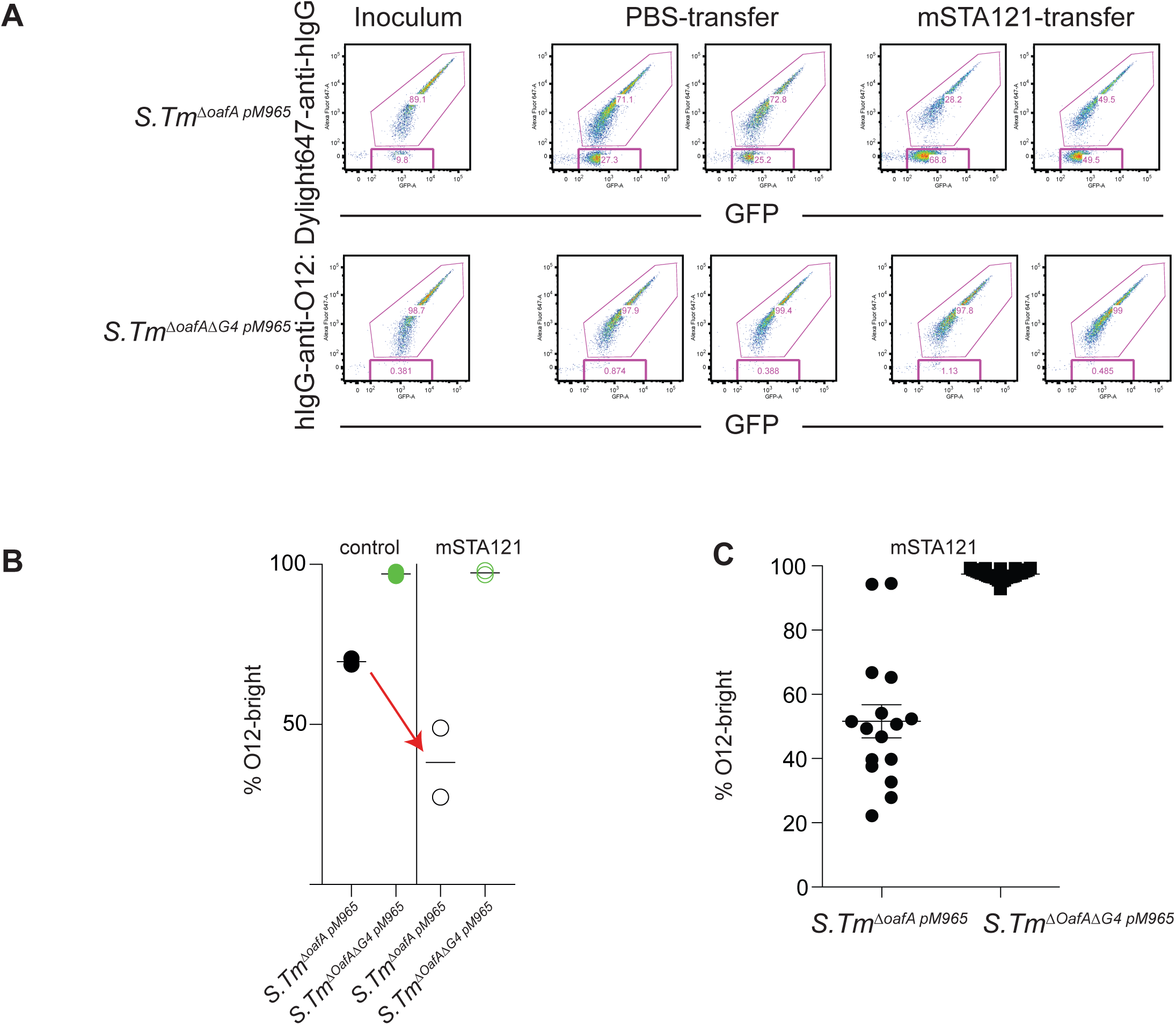
Loss of the O:12-0 epitope can be driven by adoptive transfer of O:12-0-specific IgA. C57BL/6 SPF mice received oral streptomycin to deplete the microbiota 23.5h before an intravenous injection with saline only, or with 1mg of recombinant dimeric murine IgA specific for the O:12-0 epitope (STA121). 0.5 h later all mice were orally inoculated with *S.*Tm^Δ*oafA pM965*^ or *S.*Tm^Δ*oafA*Δ*G4 pM965*^ (lacking 4 different glucosyl transferases, including *gtrC*) both carrying pM965 to drive constitutive GFP production. The adoptive transfer was repeated 12h later and all animals were euthanized at 24h post-infection. **A.** O:12-0 expression on *S.*Tm enriched from cecum content by overnight culture on 1:1000 dilution LB with selective antibiotics, determined by staining with the monoclonal antibody STA5. Flow cytometry plots shown have been gated on scatter only – see Fig. S1 for example. **B.** Quantification of the O:12-0-high fraction of *S.*Tm from A. **C.** Individual clones of *S.*Tm of the indicated genotype were recovered from the cecal content of mice from A that had received an adoptive transfer of mSTA121 and individual clones, cultured overnight in LB were analysed as in A and B for fraction of O:12-0-high cells.

**Fig. ED5:**
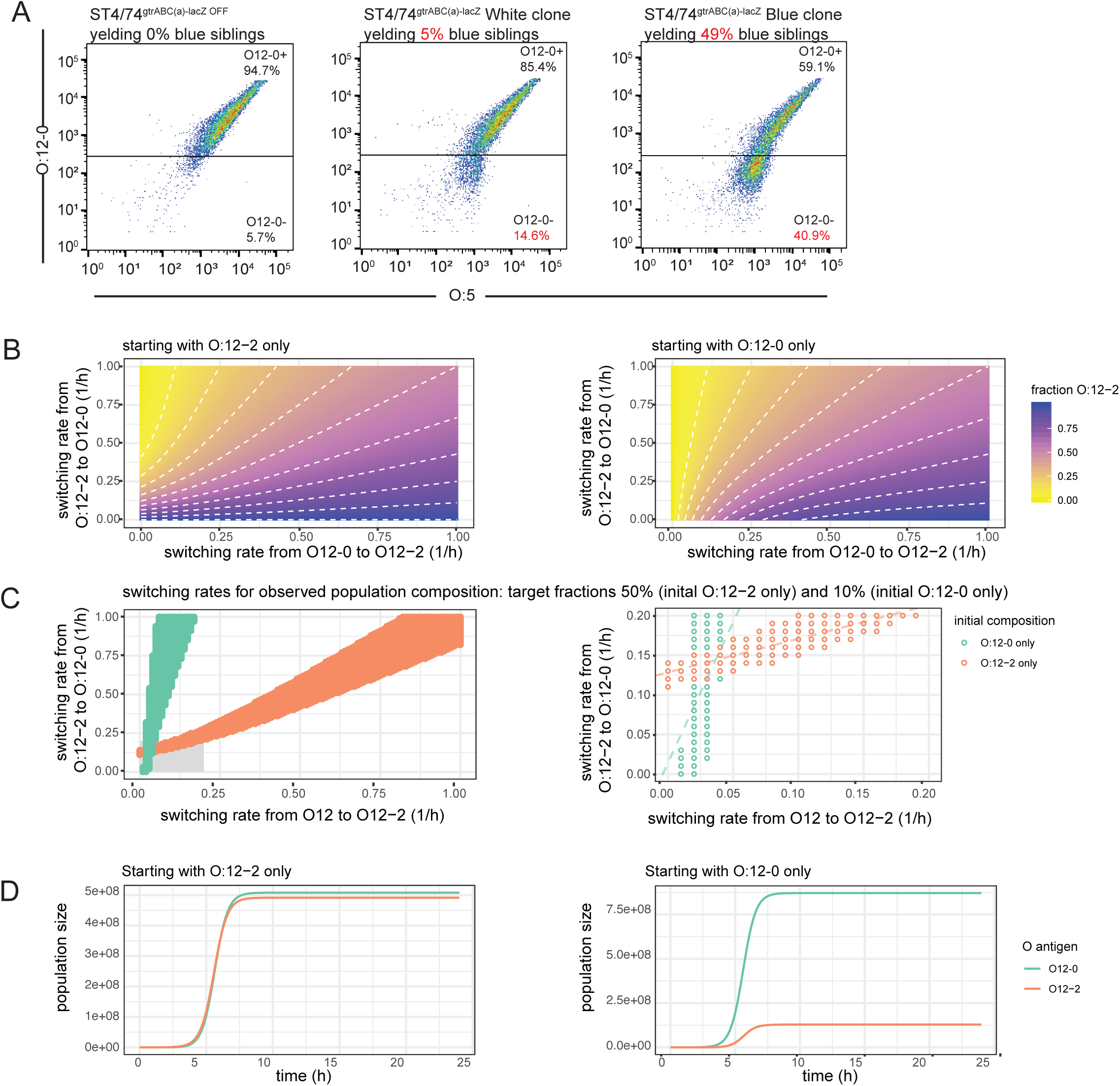
Phase-variation and selection, without a shift in switching rate, underly recovery of O:12-2 producing clones from vaccinated mice. **A.** Comparison of fractions of O:12-0-positive and O:12-0-negative bacteria (in fact O:12-2) determined by flow cytometry (gating – see Fig.S1) staining with typing sera and by blue-white colony counts using a *gtrABC*-*lacZ* reporter strain and overnight cultures from individual clonal colonies. **B-D:** Results of a mathematical model simulating bacterial growth and antigen switching (see supplementary methods). **B.** Switching rates from O:12-0 to O:12-2 and from O:12-2 to O:12-0 were varied computationally, and the fraction of O:12-2 was plotted after 16 h of growth. Left-hand plot depicts the results of the deterministic model when starting with 100% O:12-2, right-hand plot depicts the results when starting with 100% O:12-0. **C.** depicts only the switching rates that comply with the experimentally observed antigen ratios after overnight growth (90% O:12-0 when starting with O:12-0, and 50% O:12-0 when starting with O:12-2). Right-hand plot is a zoomed version showing values for switching rates between 0 – 0.2 h^-1^ (marked by a grey rectangle). Dashed lines are linear regressions on the values in this range, and their intersection marks the switching rates used for the stochastic simulation in (D). **D**. Simulation results of bacterial population growth, when starting with only O:12-2 (left-hand plot) or only O:12-0 (right-hand plot). *µ* = 2.05*h*^-1^ was kept constant in all simulations; switching rates were kept constant at *s*_->12-0_ = 0.144*h*^-1^ and *s*_->12-2_ = 0.0365*h*^-1^; the starting populations were always individuals of the indicated phenotype; carrying capacity was always *K* = 10^9^ cells. Time resolution for the simulations is 0.2*h*.

**Fig. ED6:**
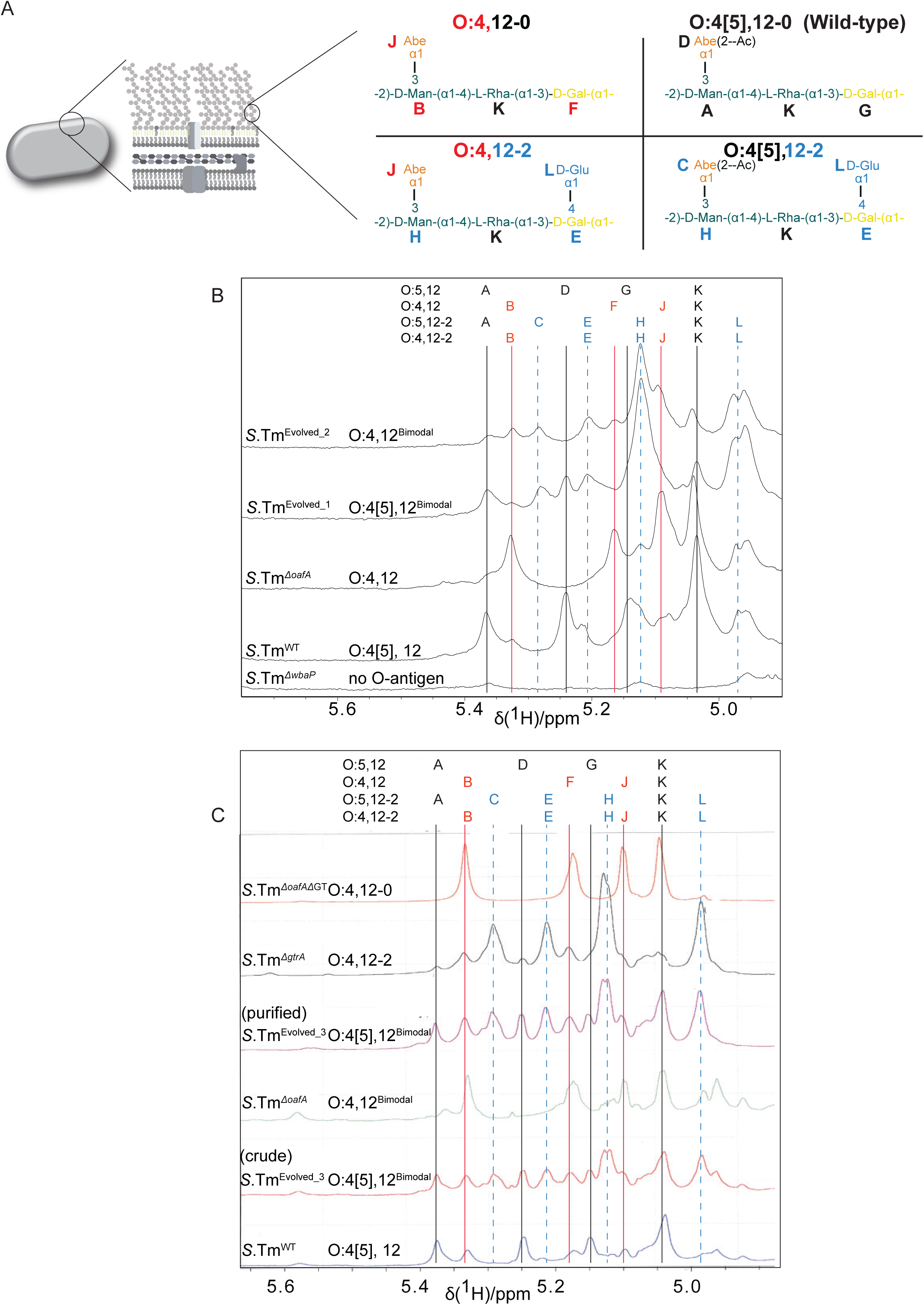
NMR of purified LPS and HR-MAS ^1^H-NMR confirms O-antigen structures in evolved clones. **A.** Schematic diagram of expected NMR peaks for each molecular species **B.** HR-MAS ^1^H-NMR spectra. Spectra show predicted peak positions and observed spectra for C1 protons of the O-antigen sugars. **C.** ^1^H NMR of purified LPS from the indicated strains. Note that non-acetylated abequose can be observed in wild type strains due to spontaneous deacetylation at low pH in late stationary phase cultures^54^. A *gtrA* mutant strain is used here to over-represent the O:12-2 O-antigen variant due to loss of regulation^5^.

**Fig ED7:**
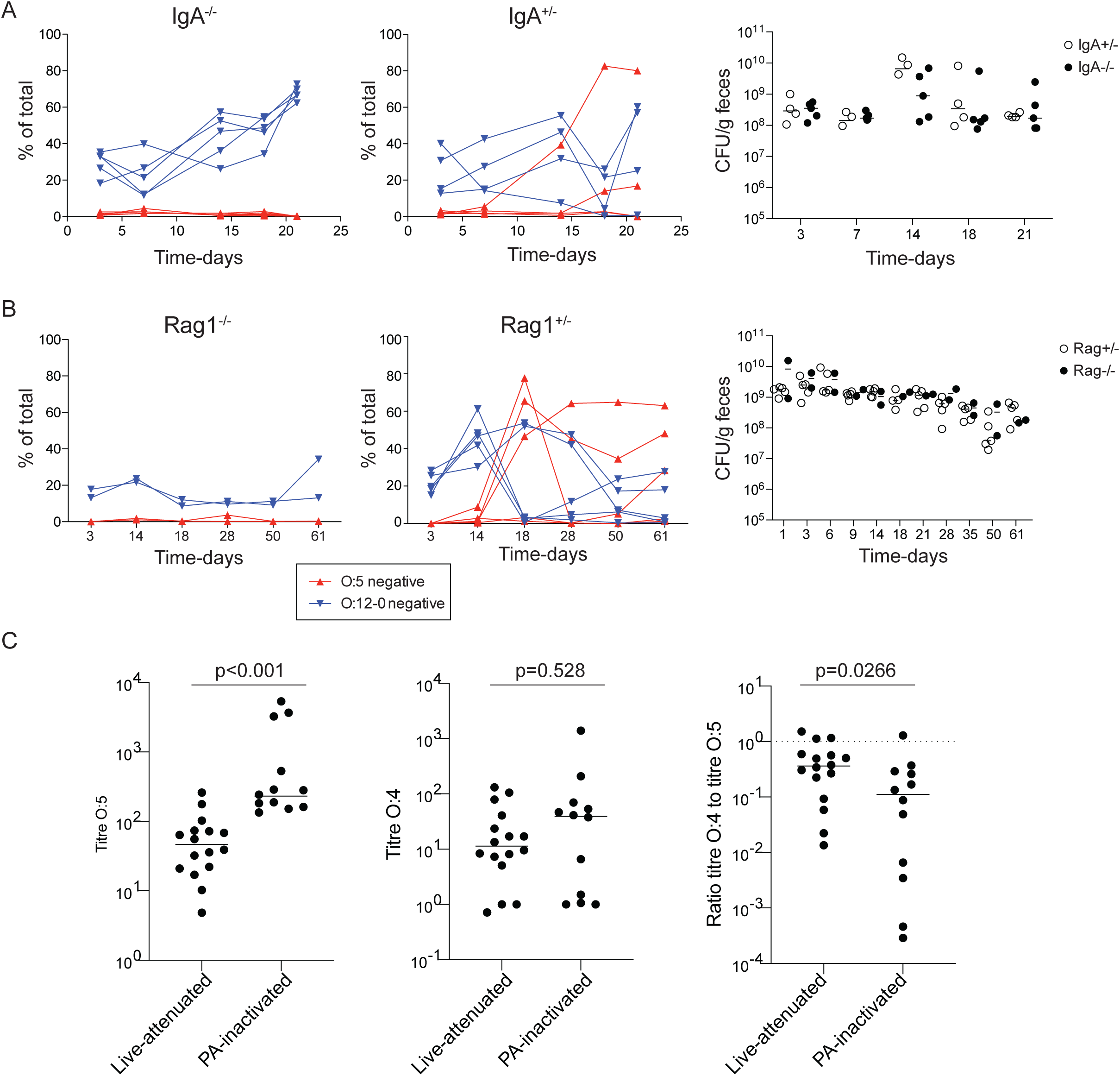
*S*.Tm O-antigen variants arise during chronic *S.*Tm infections, dependent on a specific IgA response. IgA^-/-^ **(A**) and Rag1^-/-^ **(B)** and heterozygote littermate controls (C57BL/6-background) were pre-treated with streptomycin and infected with *S*.Tm^Δ*sseD*^ orally. Fecal *S.*Tm were enriched overnight by culturing a 1:2500 dilution of feces in LB plus kanamycin. These enrichment cultures were then stained for O:5 and O:12-0 and analysed by flow cytometry (gating as in Fig. S1-4). The fraction of the population that lost O:5 and O:12-0 antisera staining is shown over time, as well as the total CFU/g in feces. Both immunocompetent mouse strains show increased O:5-negative *S.*Tm in the fecal enrichments from day 14 post-infection: approximately when we expect to see a robust secretory IgA response developing. These changes are not observed in Rag1-deficient or IgA-deficient mice. The kinetics of O:5-loss are likely influenced by development or broader IgA responses as the chronic infection proceeds. *Note that lines joining the points are to permit tracking of individual animals through the data set, and may not be representative of what occurs between the measured time-points.* **C**. Titres of intestinal lavage IgA specific for O:4[5] (*S.*Tm^wt^,O:4[5], 12-0) and O:4(*S.*Tm^Δ*oafA*^,O:4,12-0), presented as the dilution of intestinal lavage required to give an IgA-staining MFI=1000 by bacterial flow cytometry, and the ratios of these titres. Samples: d28 post-vaccination with PA-STm^wt^ (n=12) or d35 post-colonization with live-attenuated *S.*Tm (n=8 *S*.Tm^Δ*aroA*^ + n=8 *S*.Tm^Δ*sseD*^), This revealed a weaker, but less biased IgA response in mice infected with the live-vaccine strain, when compared to that induced by the inactivated oral vaccine. Results of 2-tailed Mann-Whitney U tests shown.

**Fig. ED8:**
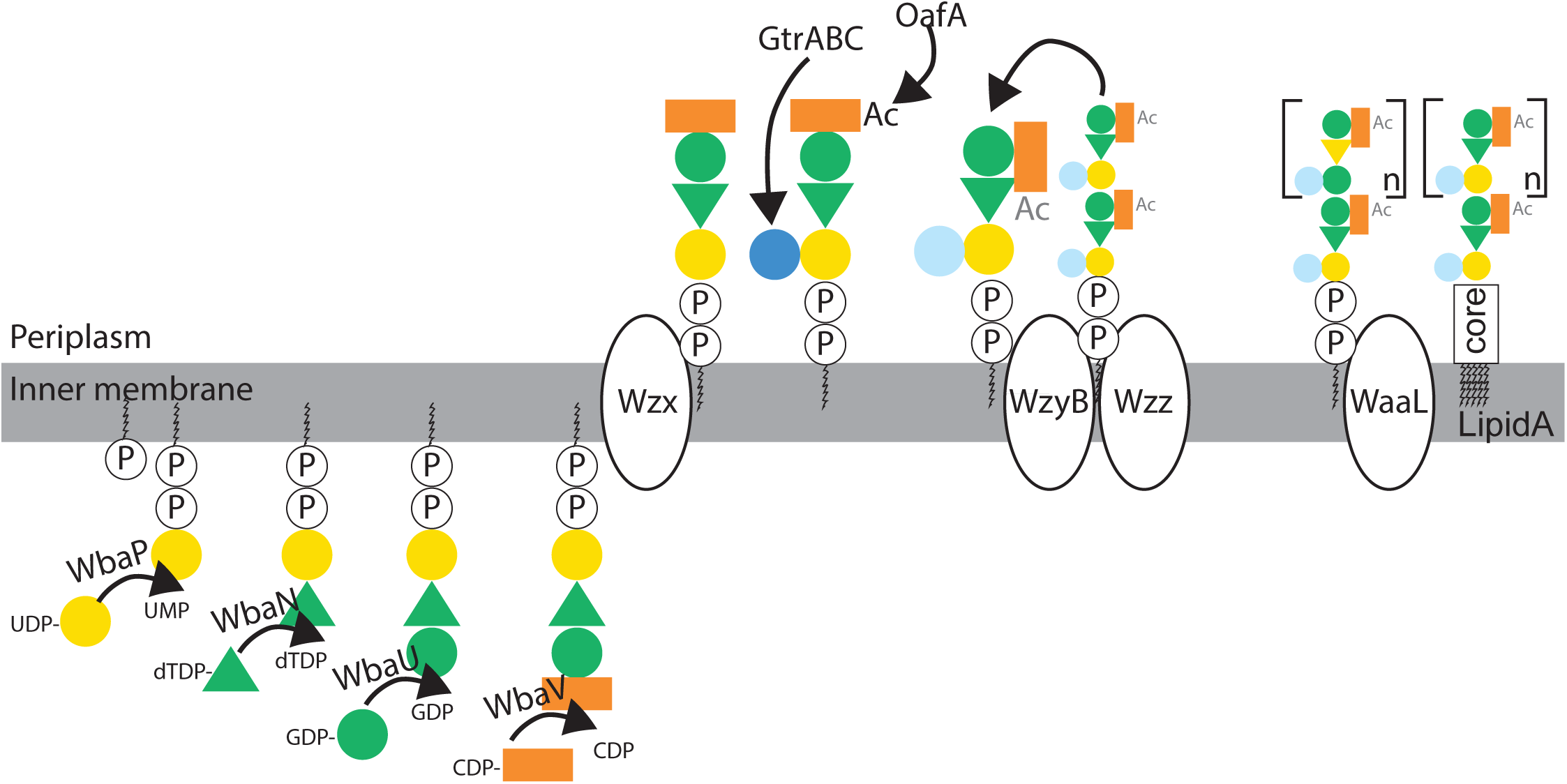
Schematic of *S.*Tm O-antigen synthesis (based on^68^)

**Fig. ED9:**
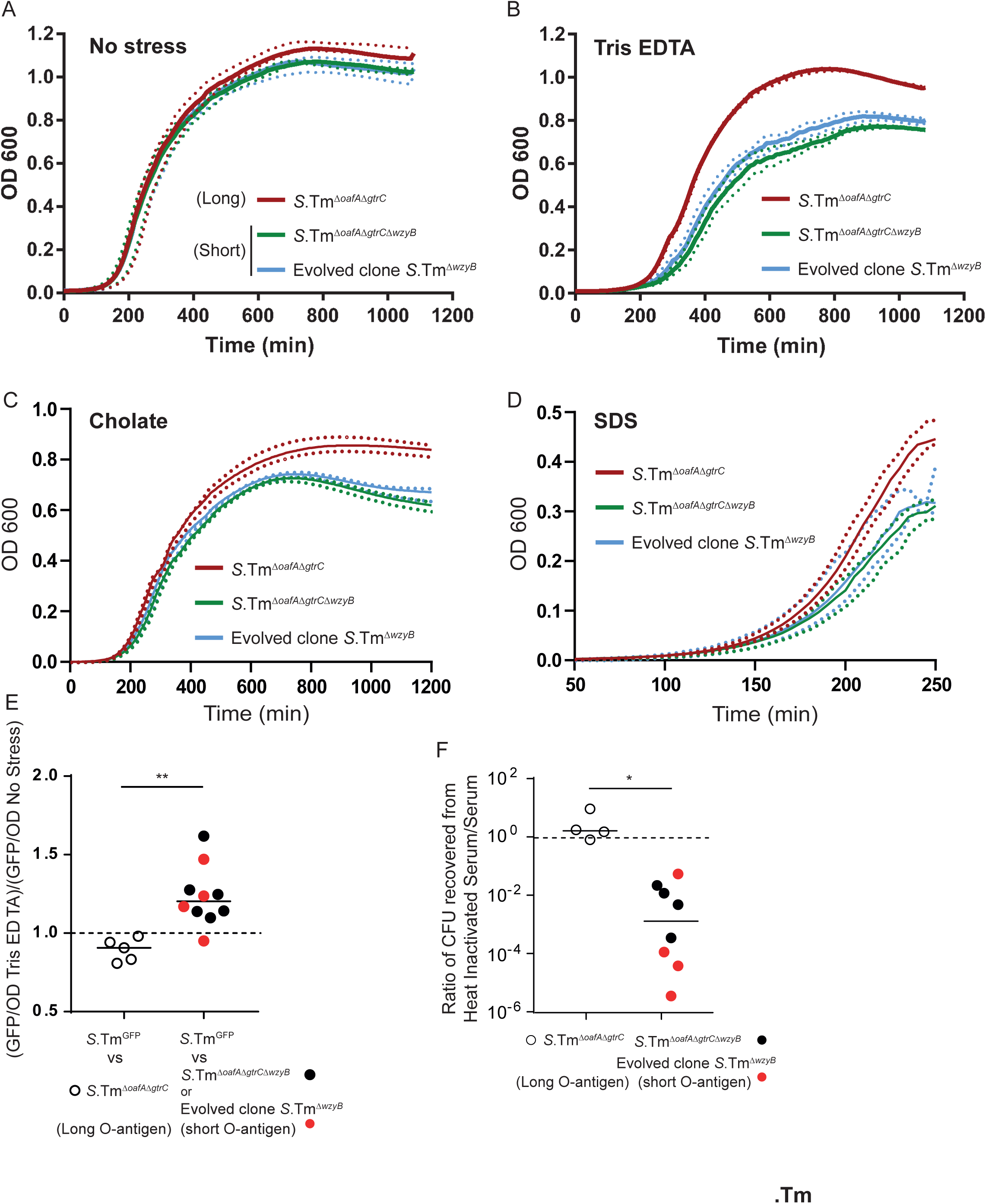
Synthetic and natural deletions of *wzyB* reduce the fitness of *S*.Tm in presence of Tris-EDTA, Cholate, SDS and serum complement. The deletion of *wzyB* does not affect the growth of *S*.Tm or *S*.Tm^Δ*oafA* Δ*gtrC*^ in LB (No stress) **(A)** but impairs growth in presence of Tris-EDTA **(B)**, 2% cholate **(C)** and 0.05% SDS **(D)**. Dashed lines represent the range of variations between the n=4 pooled experiments. (**E)**. Relative fitness of the long versus short O-antigen in the presence of membrane stress as quantified by competitive growth of *S*.Tm^GFP^ against *S*.Tm^Δ*oafA* Δ*gtrC*^, *S*.Tm ^Δ*oafA* Δ*gtrC* Δ*wzyB*^ or an evolved *S*.Tm^Δ*wzyB*^, in LB with or without Tris-EDTA. 2-tailed Mann-Whitney U test. ** p=0.0013 **(F)** Loss of complement resistance in evolved and synthetic *wzyB* mutants revealed by relative CFU recovery after treatment with heat-inactivated and fresh human serum. Mann-Whitney U 2-tailed tests * p=0.0167

**Fig. ED10:**
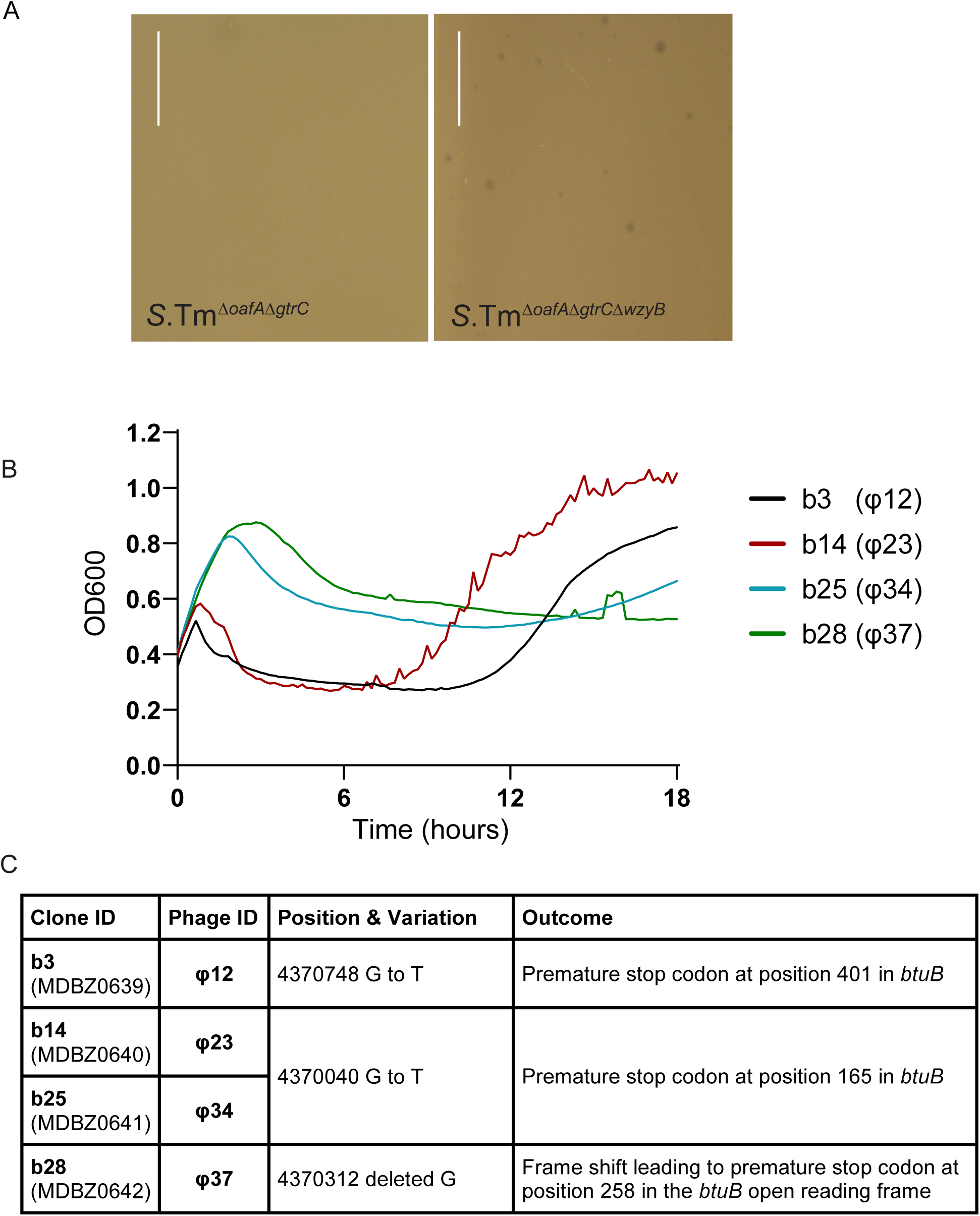
Analysis of bacteriophages preferentially infecting short O-antigen *S*. Tm mutants. **A.** Lysis plaques observed on lawns of *S*.Tm *^ΔgtrC ΔoafA^* and *S*.Tm *^ΔgtrC ΔoafA ΔwzyB^* isogenic mutants exposed to wastewater samples. Scale = 1cm. This phenocopies the observation with naturally arising *wzyB* mutants **B.** Growth curves of *S*.Tm *^ΔgtrC ΔoafA ΔwzyB^* exposed to purified bacteriophages from Fig. 4D. The re-growing *S.*Tm clones were isolated for sequencing. The mutations identified and their effects are listed in the table below (**C**), confirming *btuB* as the most likely exposed outer-membrane receptor for these phages.

## Supplementary Materials

### Supplementary tables and movies

**Table S1:** Strains and plasmids used in this study^4, 43, 45, 50, 69–71^

**Table S2:** Details of primers used in strain construction, testing and sequencing

**Table S3:** Details of mutations found in resequenced O12-0 or O12 bimodal evolved clones studied by REC-Seq as shown in Fig. 2D-G. Numbers indicate the position of the mutation, numbers in brackets indicates the percentage of reads were the mutation was detected.

**Table S4:** Details of experiments where *S.*Tm evolution was tracked, as well as further information on mice used and on clones analysed.

### Supplementary Movies A and B

Visualization of O:12 phase variation using live-cell immunofluorescence. Cells expressing GFP (green) pre-stained with fluorescently-labeled recombinant murine IgA specific for the O:12-0 epitope (red) were loaded into a microfluidic chip for time-lapse microscopy. Cells were fed continuously *S*.Tm-conditioned LB containing fluorescently-labeled recombinant murine IgA STA121 specific for the O:12-0 epitope.

(A) Loss and (B) gain of antibody reactivity (red staining) was observed, indicative of O:12 phase variation.

### Supplementary Figures

**Fig. S1:**
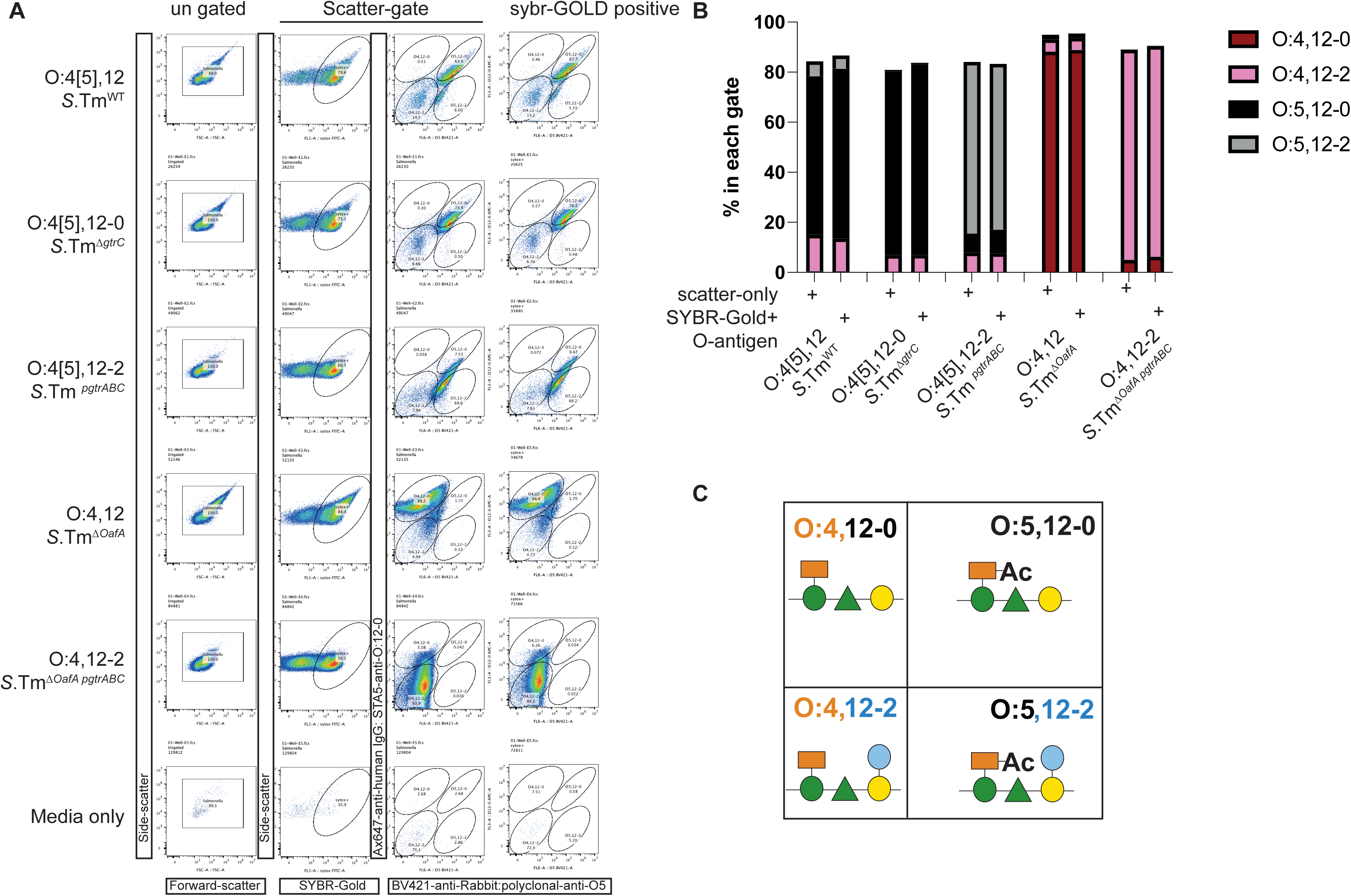
Difco Rabbit-polyclonal anti-O:5 and human monoclonal STA5 (specific for O:12-0) can be used to distinguish *Salmonella* with known O-antigen type, and can be distinguished from contaminants without DNA dyes. Overnight cultures of the indicated *S.* Typhimurium strains we made in 0.2µm-filtered LB containing streptomycin (50µg) or Ampicillin (100µg/ml to select for plasmid-maintenance of pgtrABC-containing strains). 1µl of an OD_600_=2 culture was stained in 0.2µm-filtered PBS/0.05%Azide with 1:200 Rabbit polyclonal anti-O:5 and 6µg/ml STA5. Brilliant-violet-421-Donkey-anti-Rabbit (Biolegend) and Alexa-647-anti-human IgG (Jacksom Immunolabs) were used at a 1:200 dilution, and SybrGold at 1:10’000 dilution. Samples were acquired on a Beckmann Coulter Cytoflex-S. **A.** Full gating is shown for each sample and the final analysis of O:5 versus O:12-0 staining is shown both for the entire population gated on Forward- and Side-scatter or only on DNA-dye-positive *Salmonella.* Note that the live bacteria do not stain uniformly with. SybrGold. **B.** Quantification of the O-antigen variant distribution within each strain, when gating on the entire population of the SybrGold-positive fraction reveals no difference in the analysis when DNA dyes are omitted. A sample of LB cultured overnight and treated as the samples and acquired for the same length of time as the samples (“Media only”) reveals very little background noise in our flow cytometry analysis. **C.** Schematic diagram of the O-antigen structures present on bacteria in each quadrant on the scatter plots.

**Fig. S2:**
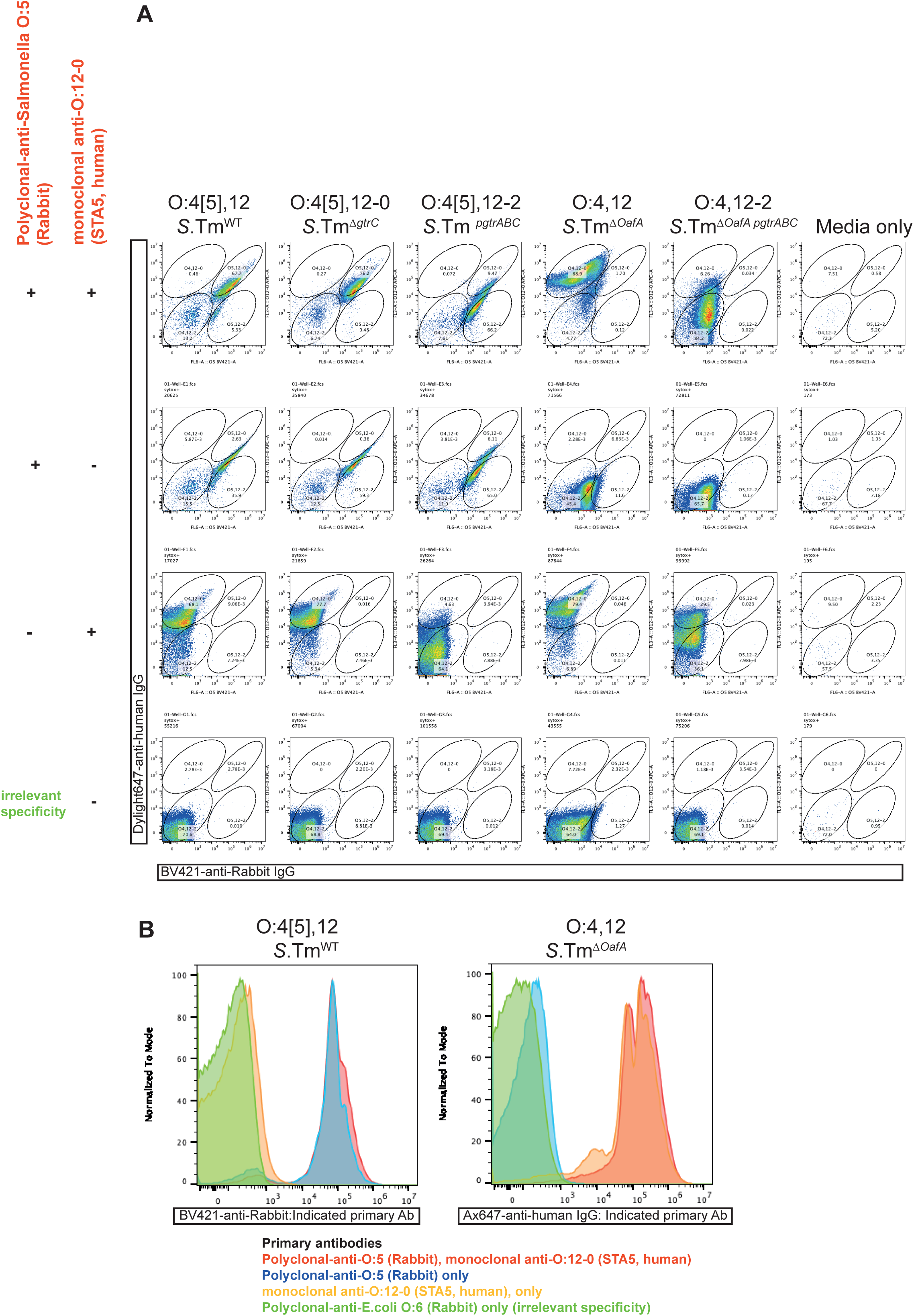
Controls for the specificity of Rabbit-polyclonal-anti-O:5 and STA5 staining. Overnight cultures of the indicated *S.* Typhimurium strains we made in 0.2µm-filtered LB containing streptomycin (50µg) or Ampicillin (100µg/ml to select for plasmid-maintenance of pgtrABC-containing strains). 1µl of an OD_600_=2 culture was stained in 0.2µm-filtered PBS/0.05%Azide with the indicated combinations of 1:200 Rabbit polyclonal anti-O:5, 1:200 Rabbit polyclonal anti-E.coli O:6, 6µg/ml STA5. Brilliant-violet-421-Donkey-anti-Rabbit (Biolegend) and Alexa-647-anti-human IgG (Jacksom Immunolabs) were used at a 1:200 dilution as secondary reagents in all stainings. Samples were acquired on a Beckmann Coulter Cytoflex-S. **A.** Samples were gated on Forward- and Side-scatter as in Fig. S1. This reveals good specificity of the antibodies with the exception of low level cross-reactivity of the anti-human IgG for the rabbit polyclonal antibody. However, the background generated by this staining is much lower than the real positive signal and does not alter interpretation of our data. **B.** Representative histogram overlays of the above stainings indicating antibody specificity.

**Fig. S3:**
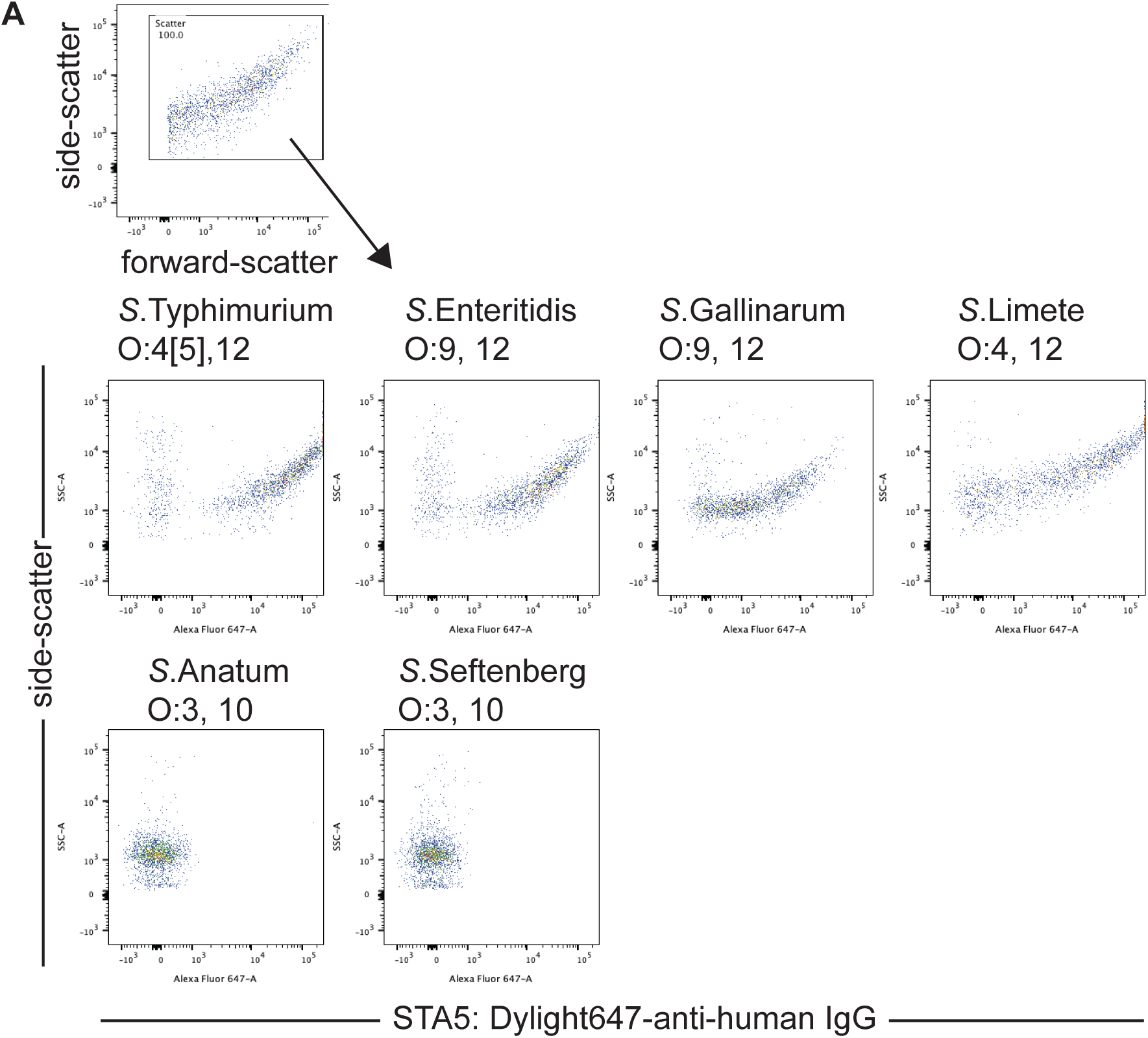
Characterization of the specificity of STA5 using diverse *Salmonella enterica* serovars and recombinant *S*.Tm strains. **A.** Recombinant monoclonal STA5 human IgG1 was used to surface stain overnight cultures of the indicated *Salmonella enterica* serovars. Bacterial surface binding was detected with a Dylight-647-anti human IgG secondary antibody and analysed by flow cytometry. STA5 binds to all serovars that include the O:12 epitope. Top panel shows the pre-gating strategy, which served only to remove events landing on the axes in forward-scatted and side-scatter measurements

**Fig. S4:**
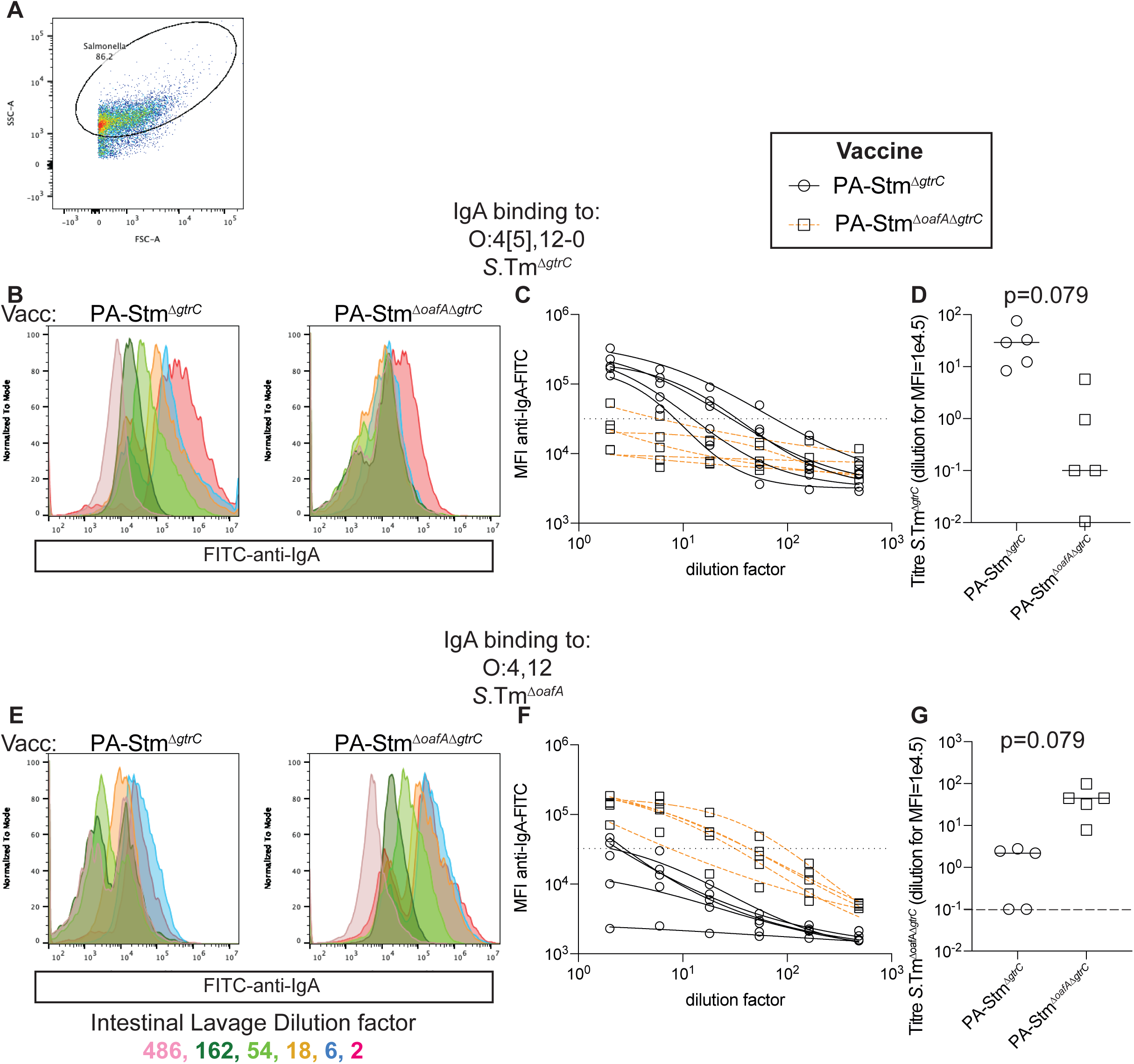
Raw data for intestinal IgA titre calculations shown in Fig. 1F and G (binding to *S.*Tm. ^Δ*gtrC*^ (O:4[5],12-0) and ***S.*Tm**^Δ*oafA* Δ*gtrC*^ (O:4,12-0). **A.** Forward- and side-scatter plot showing gating based on scatter, used for all analysis. **B.** Representative overlayed histograms of *S.*Tm ^Δ*gtrC*^ stained with intestinal lavage from a C57BL/6 mouse orally vaccinated once per week for 4 weeks with PA-STm^Δ*gtrC*^ (left) and PA-STm^Δ*oafA* Δ*gtrC*^ (right). BV421-conjugated anti-mouse IgA was used as a secondary antibody to reveal IgA coating of *S.*Tm. Colours represent different dilutions of the intestinal lavages ranging from a dilution factor of 2 (red) to 486 (pink). **C.** Intestinal lavage dilution factor plotted against the median fluorescence intensity of IgA staining (circles: PA-STm^Δ*gtrC*^-vaccinated, squares: PA-STm^Δ*gtrC*^-vaccinated) for all mice shown in Fig. 1F and G. Lines (black = PA-STm^Δ*gtrC*^-vaccinated, orange = PA-STm^ΔoafAΔ*gtrC*^-vaccinated) indicate 4-parameter logisitic curves fitted to these values using least-squares non-linear regression. **D**. Titres calculated from the fitted curves as the intestinal lavage dilution giving a median fluorescence intensity of staining = 1000 for each curve shown in C. Line indicates median value. P value of 2-tailed Mann-Whitney U test. **E.** Representative overlayed histograms of *S.*Tm ^Δ*oafA*Δ*gtrC*^ stained with intestinal lavage from a mouse orally vaccinated once per week for 4 weeks with PA-STm^Δ*gtrC*^ (left) and PA-STm^Δ*oafA* Δ*gtrC*^ (right). BV421-conjugated anti-mouse IgA was used as a secondary antibody to reveal IgA coating of *S.*Tm. Colours represent different dilutions of the intestinal lavages ranging from a dilution factor of 2 (red) to 486 (pink). **F.** Intestinal lavage dilution factor plotted against the median fluorescence intensity of IgA staining (circles: PA-STm^Δ*gtrC*^-vaccinated, squares: PA-STm^Δ*gtrC*^-vaccinated) for all mice shown in Fig. 1F and G. Lines (black = PA-STm^Δ*gtrC*^-vaccinated, orange = PA-STm^ΔoafAΔ*gtrC*^ -vaccinated) indicate 4-parameter logisitic curves fitted to these values using least-squares non-linear regression. **G**. Titres calculated from the fitted curves as the intestinal lavage dilution giving a median fluorescence intensity of staining = 1000 for each curve shown in F. Line indicates median value. P value of 2-tailed Mann-Whitney U test. All vaccinated mice were C57BL/6 and had an SPF microbiota. Note the significantly higher titres of IgA specific for the vaccination strain than the mis-matched strain.

**Fig. S5:**
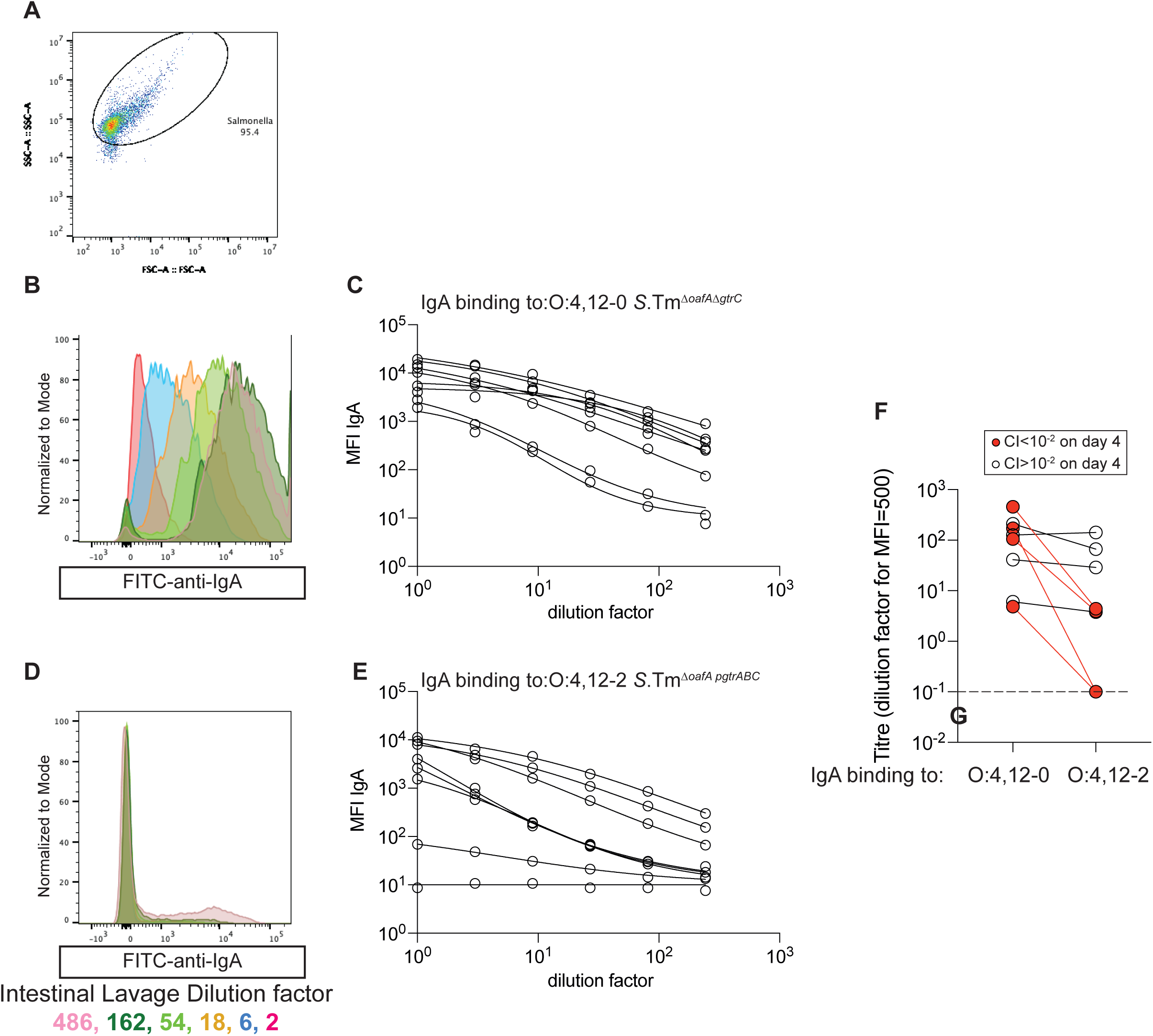
Raw data for intestinal IgA titre calculations shown in Fig. 1H (binding to *S.*Tm ^Δ*oafA,pgtrABC*^ (O:4,12-2) and ***S.*Tm**^Δ*oafA* Δ*gtrC*^ (O:4,12-0). **A.** Forward- and side-scatter plot showing gating based on scatter, used for all analysis. **B.** Representative overlayed histograms of *S.*Tm^Δ*oafA* Δ*gtrC*^ stained with intestinal lavage from a mouse orally vaccinated once per week for 4 weeks with PA-S Tm^Δ*oafA* Δ*gtrC*^. BV421-conjugated anti-mouse IgA was used as a secondary antibody to reveal IgA coating of *S.*Tm. Colours represent different dilutions of the intestinal lavages ranging from a dilution factor of 2 (red) to 486 (pink). **C.** Intestinal lavage dilution factor plotted against the median fluorescence intensity of IgA staining for all mice shown in Fig. 1H. Lines indicate 4-parameter logisitic curves fitted to these values using least-squares non-linear regression. **D**. Representative overlayed histograms of *S.*Tm^Δ*oafA pgtrABC*^ stained with intestinal lavage from a mouse orally vaccinated once per week for 4 weeks with PA-S Tm^Δ*oafA* Δ*gtrC*^. BV421-conjugated anti-mouse IgA was used as a secondary antibody to reveal IgA coating of *S.*Tm. Colours represent different dilutions of the intestinal lavages ranging from a dilution factor of 2 (red) to 486 (pink). **F.** Intestinal IgA Titres calculated from the fitted curves as the intestinal lavage dilution giving a median fluorescence intensity of staining = 1000 for each curve shown in E and C. Lines link the same lavage titred against *S.*Tm^Δ*oafA* Δ*gtrC*^ and *S.*Tm^Δ*oafA pgtrABC*^. Red symbols and lines correspond to samples in which a strain able to phase-vary O:12 out-completed the O:12-0-locked strain by more than 100-fold on day 4. In each of these mice, the IgA titre specific for *S.*Tm with an O:12-0 epitope was higher than the titre of IgA specific for the phase-varied O:12-2 variant, whereas in mice where the ability to phase-vary O:12 did not confer a selective advantage, titres against *S.*Tm^Δ*oafA* Δ*gtrC*^ and *S.*Tm^Δ*oafA pgtrABC*^ were similar. All vaccinated mice were C57BL/6 and had an SPF microbiota.

**Fig. S6:**
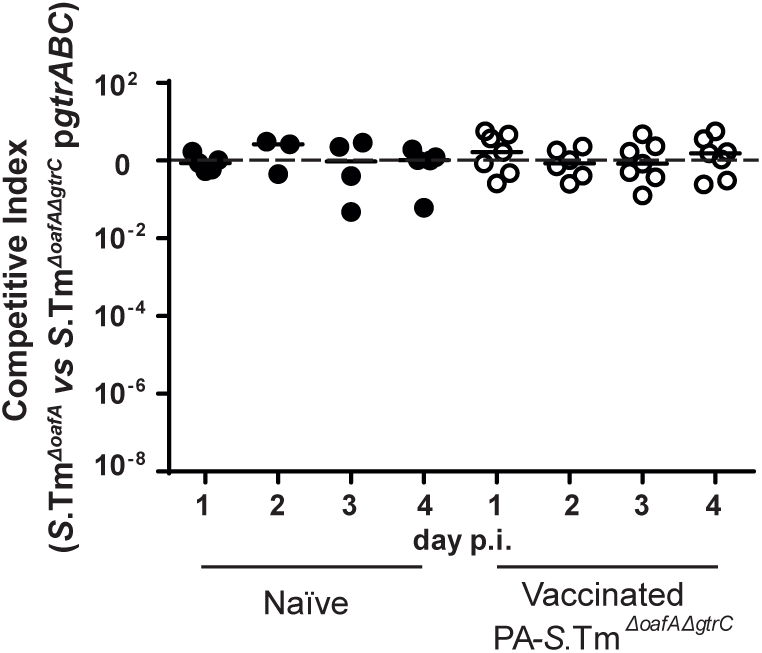
The Δ*gtrC* mutation can be complemented in trans: C57BL/6 mice were vaccinated and pre-treated as in Fig. 1. The inoculum contained a 1:1 ratio of *S.*Tm^Δ*oafA*^ and *S.*Tm^Δ*oafA* Δ*gtrC*^ p*gtrABC*. Competitive index in feces was determined by differential selective plating over 4 days post-infection.

**Fig. S7:**
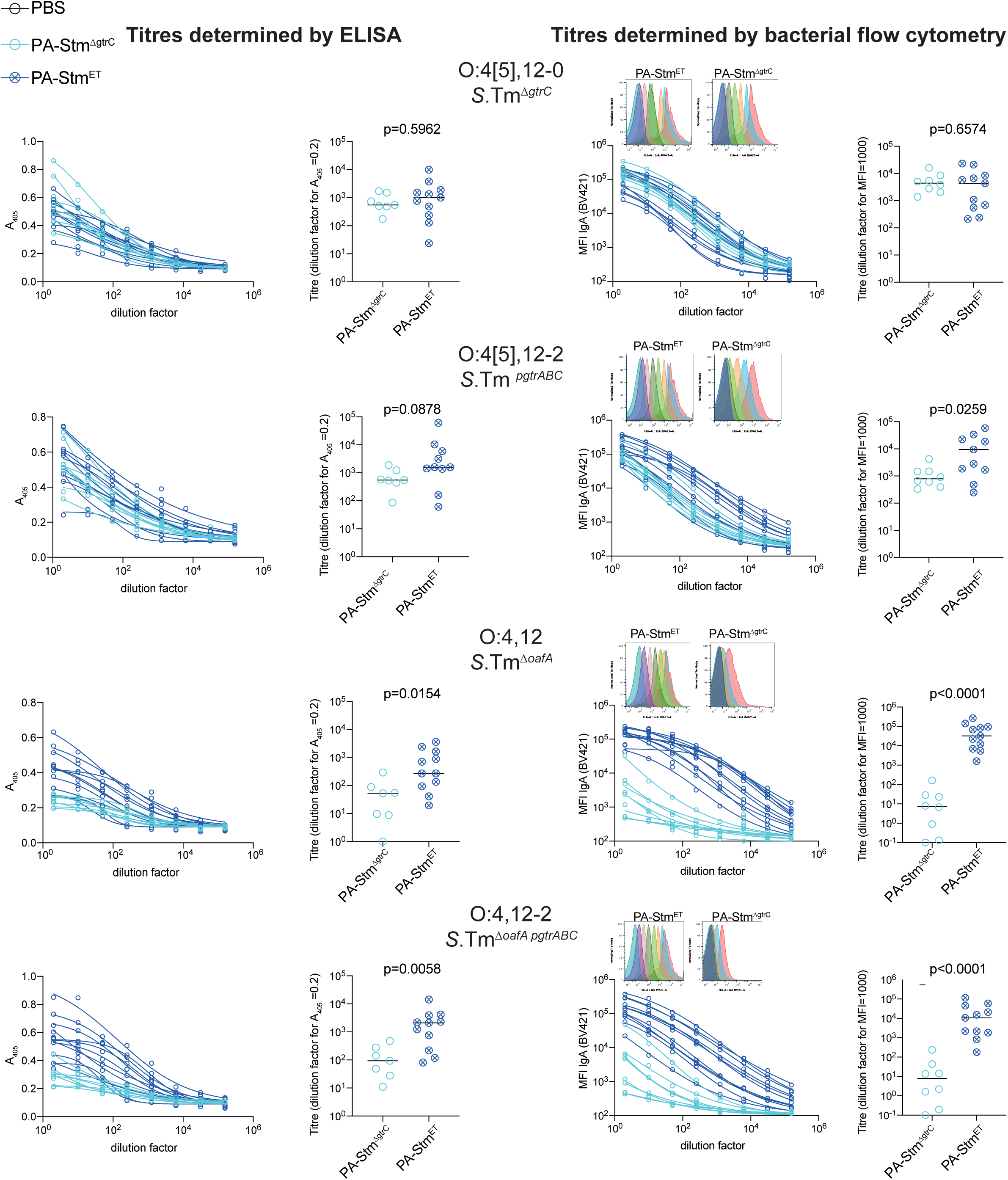
Intestinal lavage IgA titre calculations for uninfected C57BL/6 mice vaccinated with PA-STm^Δ*gtrC*^ and PA-STm^ET^ by dirty-plate ELISA and flow cytometry. C57BL/6 mice received either PA-STm^Δ*gtrC*^ (n=8) and PA-STm^ET^ (n=11) per os once per week for 4 weeks. On d28, mice were euthanized and intestinal lavages collected and cleared by centrifugation. Overnight cultures of *S.*Tm^Δ*gtrC*^, *S.*Tm*^pgtrABC^ S.*Tm^Δ*oafA*^ *and S.*Tm^Δ*oafA pgtrABC*^ were made in 0.2µm-filtered LB containing the relevant antibiotics. Bacteria were washed twice by centrifugation at 7000g to remove debris that may have accumulated during growth and used to coat ELISA plates (50µl of OD=1-0 per well) or as target for bacterial flow cytometry (10^5^ bacteria per sample). Titration curves plotting A405 (ELISA) or median fluorescence intensity (bacteria flow cytometry) as read-outs of IgA binding, against dilution factor of lavages were used to calculate titres from 4-parameter logistic curve-fits. Representative overlayed histograms of the flow cytometry read-out from one PA-STm^Δ*gtrC*^ and PA-STm^ET^-vaccinated mouse are shown (Colours represent different 3-fold serial dilutions of the intestinal lavage: red=2, blue=6, orange=18, green=54, dark green = 162, pink = 486. Pre-gated on scatted as in Fig. S4). P-values were calculated using 2-tailed Mann Whitney U tests. Flow cytometry and ELISA reveal similar results, but with flow cytometry giving a clearer read-out. This is likely due to binding of lysed bacterial components to the ELISA plate scaffold, including protein components that will be identical between our strains as well as antigenic, but which are not accessible on the surface of live cells, and are therefore irrelevant for protection. We have therefore used bacterial flow cytometry to titre intestinal IgA throughout the manuscript as it more straightforward to equate to IgA binding to the surface of whole, intact, live cells.

**Fig. S8:**
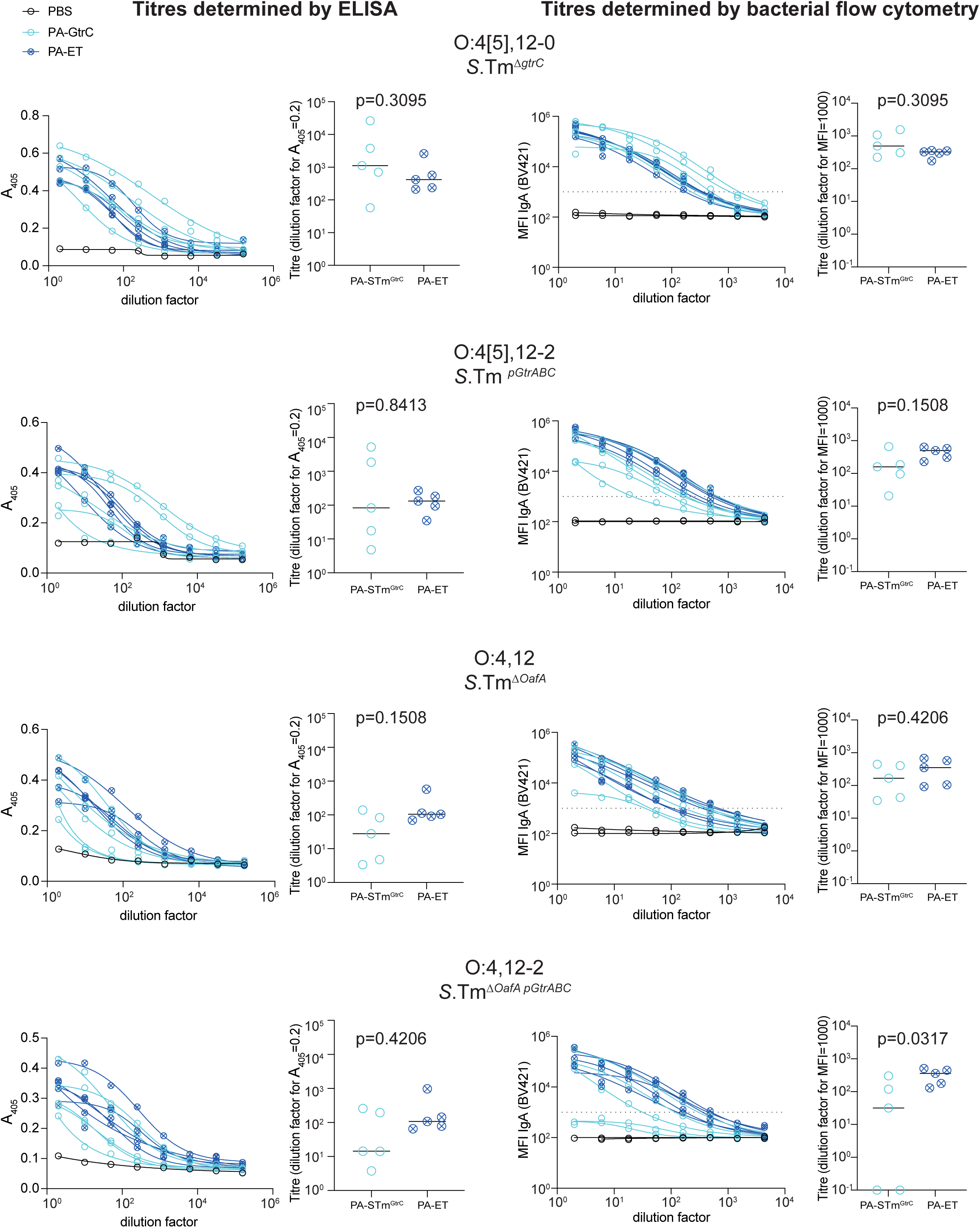
Intestinal lavage IgA titre calculations for 129S1/SvImJ mice vaccinated with PA-STm^Δ*gtrC*^ and PA-STm^ET^ and infected with *S.*Tm^WT^ for 9 days, by dirty-plate ELISA and flow cytometry. 129S1/SvImJ mice received the indicated vaccine per os once per week for 4 weeks. On d28, mice were treated with oral streptomycin and were infected with *S.*Tm^WT^. Nine days post-infection, all mice were euthanized and intestinal lavages collected and cleared by centrifugation. Overnight cultures of *S.*Tm^Δ*gtrC*^, *S.*Tm*^pgtrABC^ S.*Tm^Δ*oafA*^ *and S.*Tm^Δ*oafA pgtrABC*^ were made in 0.2µm-filtered LB containing the relevant antibiotics. Bacteria were washed twice by centrifugation at 7000g to remove debris that may have accumulated during growth and used to coat ELISA plates (50µl of OD=1-0 per well) or as target for bacterial flow cytometry (10^5^ bacteria per sample). Titration curves plotting A405 (ELISA) or median fluorescence intensity (bacteria flow cytometry) as read-outs of IgA binding, against dilution factor of lavages were used to calculate titres from 4-parameter logistic curve-fits. P-values were calculated using 2-tailed Mann Whitney U tests. Flow cytometry and ELISA reveal similar results. Note that there is some broadening of the IgA response in PA-STm^ΔgtrC^-vaccinated mice over the 9 days of infection when compared to the data in Fig. S7.

**Fig. S9:**
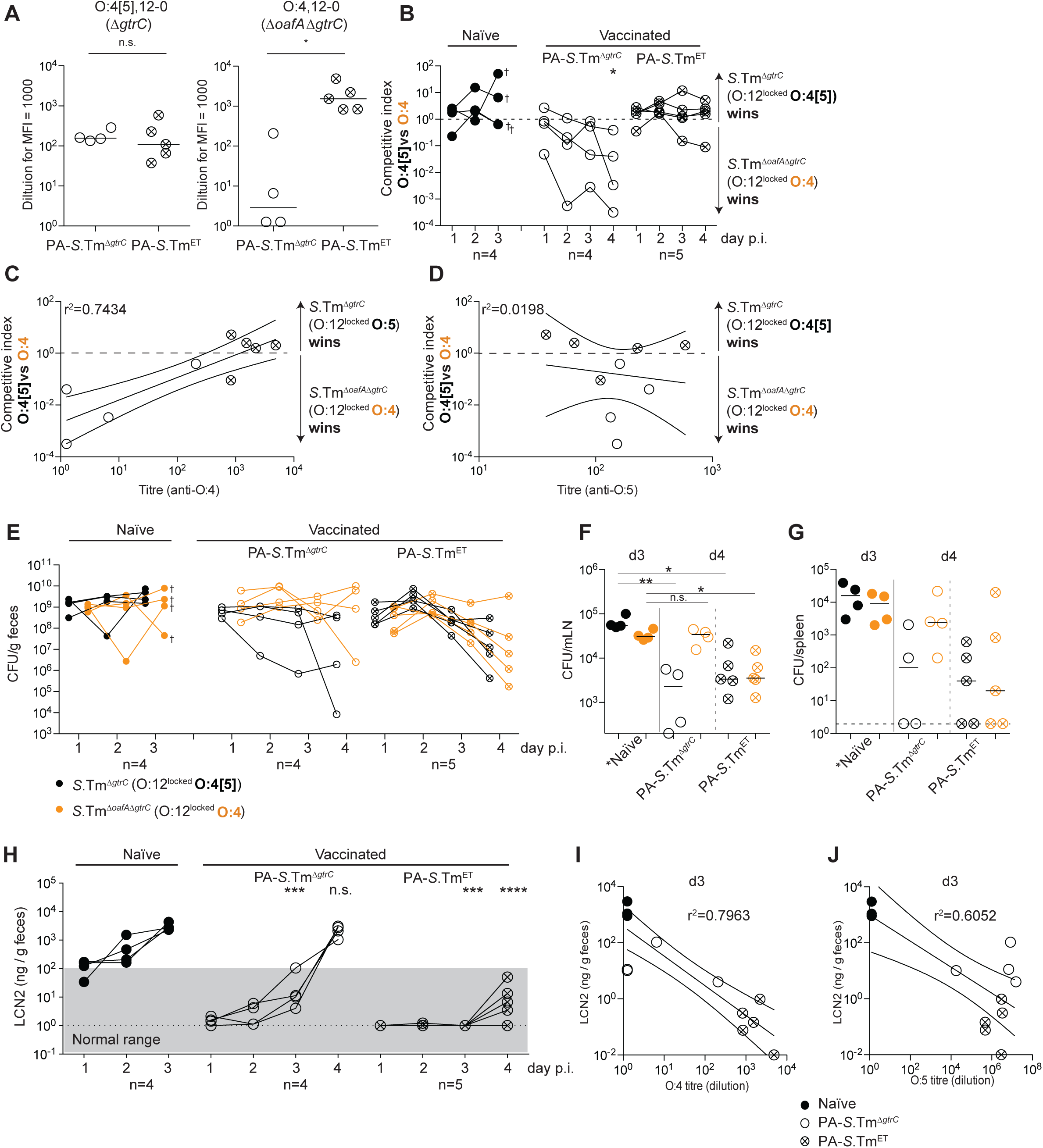
IgA-driven selective pressure functions identically in SPF Balb/c mice. **A-C**. Previous work indicated that Balb/c mice may respond better to oral vaccines and produce more secretary IgA than C57BL/6 mice (Fransen et al. 2015), therefore we tested the ability of PA-STm^ET^ to protect in Balb/c mice. Naive (closed circles), PA-*S.*Tm^Δ*gtrC*^-vaccinated (open circles) and PA-*S.*Tm^ET^-vaccinated (crossed-circles) SPF Balb/c mice were streptomycin-pretreated, infected (10^5^ CFU, 1:1 ratio of *S.*Tm^Δ*gtrC*^ and *S.*Tm^Δ*gtrC* Δ*oafA*^ per os). Note that naïve Balb/c mice were euthanized on day 3 due to severe disease. **A**. Secretory IgA titres (intestinal lavage dilution) against O:4[5], 12-0, and an O:4, 12-0 *S.*Tm. *p=0.0159 **B.** Competitive index (CFU *S.*Tm ^Δ*gtrC*^*/*CFU *S.*Tm^Δ *gtrC* Δ*oafA*^) in feces at the indicated time-points. 2-way ANOVA with Bonferroni post-tests on log-normalized values, compared to naive mice. *p<0.0285. O:4-only producing *S.*Tm outcompetes in mice vaccinated with PA-STm^Δ*gtrC*^ but not PA-STm^ET^. **C** and **D**. Correlation of the competitive index with the O:4-specific (**C**) and O:4[5]-specific (**D**) intestinal IgA titre, r^2^ values of the linear regression of log-normalized values. Open circles: Intestinal IgA from PA-*S.*Tm^Δ*gtrC*^ -vaccinated mice, crossed circles: Intestinal IgA from PA-*S.*Tm^ET^ -vaccinated mice. Lines indicate the best fit with 95% confidence interval. As both vaccinated groups have similar titres against the O:4[5]-producing S.Tm, a correlation of C.I. is observed only with the O:4-specific IgA titre **E.** CFU of *S.*Tm^Δ*gtrC*^ (black symbols) and *S.*Tm^Δ*gtrC* Δ*oafA*^ (orange symbols) per gram feces at the indicated time-points. **F and G.** CFU of *S.*Tm^Δ*gtrC*^ (black symbols) and *S.*Tm^Δ*gtrC* Δ*oafA*^ (orange symbols) per organ and day 4 post-infection (vaccinated) and day 3 post-infection (naïve). Kruskal-Wallis test with Dunn’s multiple comparison adjusted P values are shown. *p=0.022, **p=0.0085 **H.** Fecal Lipocalin 2 as a marker of inflammation in the indicated groups. 2-way repeat-measures ANOVA on log-normalized data, with Bonferroni post-tests comparing to the Naïve mice. ***p=0.0002,****p<0.0001 **I and J.** Correlation between fecal lipocalin 2 on d3 post-infection and O:4 and O:4[5]-specific intestinal IgA titres. r^2^ values of the linear regression of log-normalized values. Lines indicate the best fit with 95% confidence interval. *Note that lines joining the points in **B, E and H** are to permit tracking of individual animals through the data set, and may not be representative of what occurs between the measured time-points.* This experiment was based on the observations made in Fransen et al^72^ that better IgA-mediated protection is achieved in Balb/c mice than in C57BL/6 mice in response to live-attenuated vaccines. However, both mouse lines behave similarly in this model.

**Fig S10:**
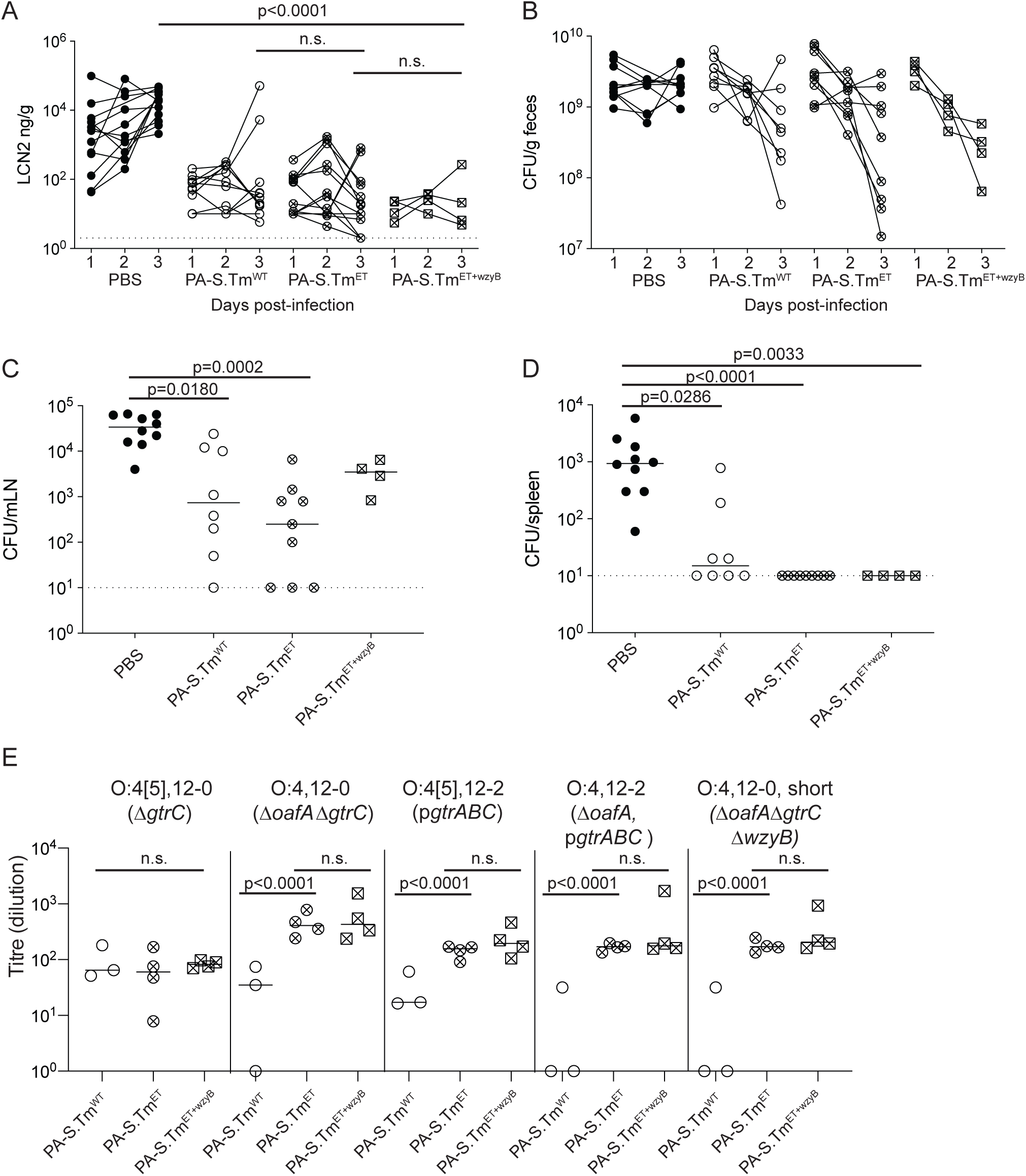
PA-STm^ET^ mediated effects are not improved by addition of *S*.Tm^Δ*wzyB*^ to the vaccine cocktail. C57BL/6 mice were vaccinated with vehicle only (Naïve, n=10), PA-S.Tm^wt^ (n=8), PA-STm^ET^ (combined PA*-S.*Tm^Δ*gtrC*^, PA-*S.*Tm^Δ*oafA* Δ*gtrC*^, PA-*S.*Tm p*gtrABC*, and PA-*S.*Tm^Δ*oafA*^ p*gtrABC,* n=9*) or* PA-STm^ET+wzyB^ (combined PA*-S.*Tm^Δ*gtrC*^, PA-*S.*Tm^Δ*oafA* Δ*gtrC*^, PA-*S.*Tm p*gtrABC*, PA-*S.*Tm^Δ*oafA*^ p*gtrABC* and PA-*S.*Tm^Δ*oafA* Δ*gtrC* ΔwzyB^, n=4). On day 28 after the first vaccination, mice were streptomycin pre-treated and challenged with 10^5^ *S.*Tm^wt^ orally. Fecal Lipocalin-2 (LCN2) at day 1-3 post-infection, (**A**) and CFU *S.*Tm^wt^ per gram feces on day 1-3 post -infection (**B**), CFU *S.*Tm^wt^ per mesenteric lymph node (MLN) at day 3 post-infection (**C**), and CFU *S.*Tm^wt^ per spleen at day 3 post-infection (**D**). A and B, 2-way repeat-measures ANOVA on log-normalized data with Bonferroni multiple comparisons-tests reveals no significant difference between the vaccinated groups at any time-point. Adjusted p values are displayed. C and D: Kruskal-Wallis analyses **with Dunn’s multiple comparisons-tests comparing all groups** were carried out for significance. Exact adjusted p values displayed. **E.** IgA titres in intestinal lavage of an experiment not included in (Fig. 3D), and additionally showing the group PA-*S.*Tm^Δ*oafA* Δ*gtrC* Δ*wzyB*^. Titres are expressed as the dilution factor of lavage required to give an MFI=1000. 2-way repeat-measures ANOVA on log-normalized data with Bonferroni multiple comparisons-tests. *Note that lines joining the points in **A and B** are to permit tracking of individual animals through the data set, and may not be representative of what occurs between the measured time-points*.

